# Deep neurobehavioral phenotyping uncovers neural fingerprints of locomotor deficits in Parkinson’s disease

**DOI:** 10.1101/2025.04.22.650006

**Authors:** Elisa L. Garulli, Timon Merk, Ghadi El Hasbani, Burçe Kabaŏglu, Rafael De Sa, Ruben Behrsing, Dennis Doll, Michael Schellenberger, Ibrahem Hanafi, Arend Vogt, Wolf-Julian Neumann, Chiara Palmisano, Yangfan Peng, Matthias Endres, Christoph Harms, Nikolaus Wenger

**Affiliations:** Department of Experimental Neurology, Charité – Universitätsmedizin Berlin, Germany; Movement Disorders Unit, Charité – Universitätsmedizin Berlin, Germany; Center for Stroke Research Berlin, Germany; DZHK (German Center for Cardiovascular Research), Germany; DZNE (German Center for Neurodegenerative Diseases), Germany; DZPG (German Center of Mental Health), Berlin, Germany; Einstein Centre for Neuroscience Berlin, Germany; Institute of Clinical Neurobiology, University Hospital Wuerzburg, Germany; Klinik und Hochschulambulanz für Neurologie, Charité – Universitätsmedizin Berlin, Germany; Department of Neurology and Neuroscience Research Center, Charité – Universitätsmedizin Berlin, Germany; Indoc Research Europe gGmbH, Mainz, Germany

**Keywords:** Parkinson, akinesia, biomarkers, neurobehavioral

## Abstract

Gait deficits present an unresolved therapeutic challenge in Parkinson’s Disease. At the behavioral level, symptoms exhibit heterogeneity, including bradykinesia and hypokinesia during cyclical limb movement, as well as sudden interruptions in the gait sequence, also known as freezing of gait. The neural activities that drive these various deficits remain largely unknown. Here, we investigated the neural correlates of gait sequence interruptions with deep neurobehavioral phenotyping. For this, we transformed kinematic trajectories and cortical oscillations into continuous time series of multimodal feature vectors. Next, we applied machine learning, combining low-dimensional embedding with supervised classification, to identify cortical oscillation features that drive gait deficits. In a rodent Parkinson’s disease model, our approach revealed that gait, akinesia, and stationary movements occupy prominently different regions in the low-dimensional embedding space. Among the predominant features separating the states, we found Hjorth complexity and mobility to modulate with the onset of akinetic episodes. Additionally, we validated our analysis approach in two Parkinson patients with freezing of gait, where neural features in STN recordings partially reflected the findings from ECoG measurements in rodents. The presented neurobehavioral phenotyping approach is translational and can easily generalize to the analysis of other complex movement disorders. Together, our results highlight specific features of neural oscillations as potential biomarkers that may support the development of adaptive closed-loop algorithms for gait therapy in PD.

## 1 Introduction

Parkinson’s Disease (PD) is a neurodegenerative disease characterized by diverse neural and behavioral deficits, ranging from emotional components to movement [1, 2]. Motor impairments include an array of symptoms such as hypo- and bradykinesia, tremor, and Freezing of Gait (FoG). FoG has been formalized as *”brief, episodic absence or marked reduction of forward progression of the feet despite the intention to walk”* [3–6]. Current PD-therapies include dopaminergic medication, eventually combined with Deep Brain Stimulation (DBS) primarily of the Subthalamic Nucleus (STN). These interventions are effective at addressing tremor and rigidity but have moderate, or no, efficacy on gait impairments [7, 8]. The limitations of pharmaceutical and open-loop DBS, led to the exploration of adaptive DBS protocols to provide enhanced treatment effects [9–11]. Effective stimulation protocols hinge on identifying suitable biomarkers that correlate with symptoms and can be monitored to guide the intervention [11, 12]. Yet, these biomarkers have remained poorly characterized for gait impairments, and the heterogeneous nature of gait symptoms has further hindered discovery.

Beta-band oscillations (13-35 Hz in humans) are a common target for adaptive DBS [12] as they have been repeatedly shown to correlate with bradykinesia [13, 14], rigidity [14] and tremor [15]. Indeed, algorithms locking to beta bursts showed promising results addressing hemibody subscores (bradykinesia, rigidity, tremor) of the Unified Parkinson’s Disease Rating Scale (UPDRS) [11, 12]. Beta phase-locked stimulation [16] have been proposed as a further closed-loop stimulation refinement. Moreover, gamma power entrainment with standard open-loop DBS has also been shown to effectively ameliorate a diverse set of motor symptoms (most prominently bradykinesia and dystonia) [9]. For gait in Parkinson’s patients, first insights have been generated on the correlation of FoG with high beta-gamma phase-amplitude coupling [17], high beta cortico-STN coherence [18] and abnormal patterns in the theta band [19]. Moreover, the combination of multiple neural markers, into a multi-input model has been proposed as a promising strategy to address complex and heterogeneous symptom manifestations at multiple scales [20, 21]. Yet, a detailed characterisation of neural oscillation features that drive gait impairments remains to be achieved.

Here we addressed the neural correlates of locomotor deficits using deep neurobehavioral phenotyping in the unilateral 6-OHDA rodent model of Parkinson’s disease. This model offers a well established platform to explore gait impairments and inform the next generation of neuromodulation therapies [22]. A typical locomotor sequence in rodents consists of *gait*, which is the cyclical limb movements that propels the animal forward [23], and can be preceded, interrupted or followed by active or inactive behaviors [24]. Active motor behaviors include grooming, sniffing, and scanning the environment - here collectively referred to as *stationary movement*. Inactive motor behaviors include freezing, a state of akinesia that can occur as an innate response to acute stressors [25], or as a pathological symptom in the parkinsonian condition [3]. In this study, we used deep neurobehavioral phenotyping to selectively analyse the neural correlates of distinct movement states in the locomotor sequence.

We computed neurobehavioral feature vectors that segment the locomotion sequence into discrete time windows. Next, we used machine-learning models to uncover the salient features that best characterize deficits in the locomotor sequences. With this strategy we identified Hjorth complexity (deviation from a sinusoid) and mobility (frequency variance) [26] as novel parameters that correlate with akinetic episodes onset. These parameters are computationally efficient and low-latency, making them attractive target candidates for closed-loop neuromodulation. Finally, we underscored the potential utility the identified neural features in rats by confirming the same pattern of modulation in STN neural recording from a Parkinson’s patient with FoG episodes.

## 2 Results

### 2.1 Basic processing of of locomotor deficits confirms established neural biomarkers in 6-OHDA rats

To analyze neurobehavioral correlates of motor deficits, we employed *neurokin*, a self-developed toolbox (https://neurokin.readthedocs.io) to integrate multimodal data. We worked with the unilateral 6-OHDA model as a standard for locomotor deficits [22]. The left Medial Forebrain Bundle (MFB) was injected with 9.6 *µ*g of 6-hydroxydopamine (6-OHDA) in N=6 Sprague Dawley rats, resulting in the degeneration of dopaminergic innervation (**Fig 1a**). We quantified the degeneration through Tyrosine Hydroxylase (TH) staining and signal intensity measurements in striatal brain sections (**Fig 1b**), the neurotoxin led to a signal intensity loss of *>*85% (mean TH signal, left: 4.78 *±*0.51 A.U., right: 30.41 *±*4.22 A.U., P *<* 0.001) (**Fig 1c**). Sham animals (N=8) underwent the same procedure, but were injected with saline. After manifestation of deficits, rats were allowed to behave freely on the runway. To investigate the neurobehavioral correlates, we collected primary motor cortex (M1) ElectroCorticoGraphy (ECoG) signals (at a sampling rate of 24 KHz) and video-recorded the experiments (at a 50 Hz frame rate). From the videos, we manually labelled the three locomotor states: i) gait, ii) stationary movement and 3) akinesia - thus establishing ground truth labels (**Fig 1d**). A representative labelling of a runway trial in a 6-OHDA lesioned rat is displayed in **Video Supplement 1**.

**Fig. 1.**
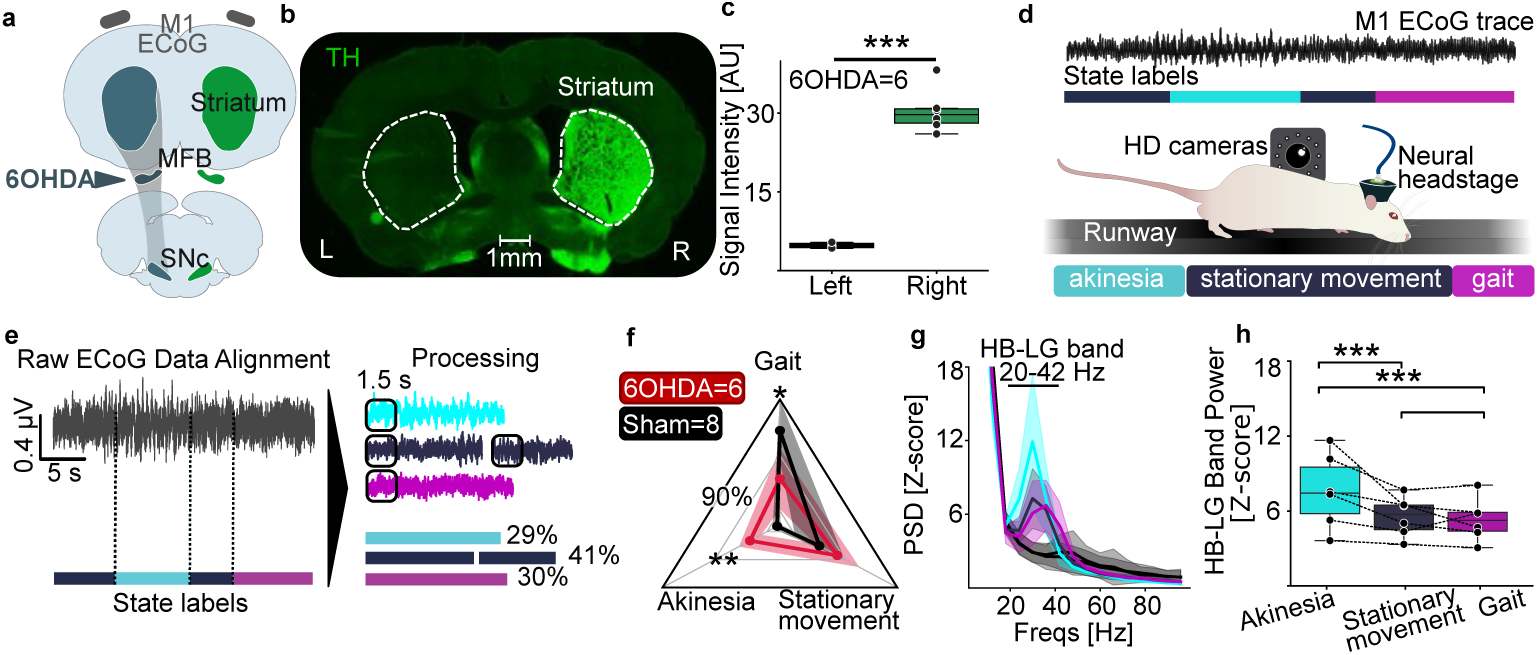
Neurobehavioral profiling of locomotor deficits in 6-OHDA rats. **a. Experimental model.** Rats were unilaterally injected with 6-OHDA in the MFB and bilaterally implanted with ECoG electrodes over the motor cortex. **b. Tyrosine-Hydroxylase (TH) stainings** highlighted successful unilateral degeneration of dopamine projections in the striatum of 6-OHDA injected rats. **c. TH-Staining: statistical quantification d. Experimental set-up.** Rats were trained to walk on a runway. Locomotor behavior was recorded from two sides with HD cameras (50 fps). Neural data from M1-ECoG were acquired with a digitizing headstage at 24 kHz. Locomotor states were visually identified during offline analysis (Nexus 2 software) **e. Modular pipeline enabled by *neurokin*.** Two *neurokin* modules (i) converted user-marked timestamps to percentage-of-trial values and (ii) aligned LFPs to compute state-specific PSDs. **f. Distribution of locomotor states.** PD rats spent more time in akinesia (P *<* 0.01) and less in gait (P *<* 0.05) than shams, (mean *±* SD). **g. State-specific PSD.** HB-LG power peaked during akinesia (cyan) versus stationary (purple), gait (magenta), and shams (black; akinesia trace omitted, n *<* 3). **h. Linear Mixed Model analysis.** HB-LG band amplitude was higher in akinesia than stationary or gait (both P *<* 0.001). States were modelled as a fixed effect and day nested in subject as a random effect.

Next, we used *neurokin* to process manual labels and synchronize the events to the neural data (**Fig 1e**). We then built a pipeline to compute the percentage of time spent in each state. The data showed that rats injected with 6-OHDA spent more time in akinetic states (P *<* 0.01, t-test with Bonferroni-Holm correction) and less time in gait than sham rats (P *<* 0.05, t-test with Bonferroni-Holm correction) (**Fig 1f**). Next, we calculated the corresponding Power Spectral Density (PSD) between 5 and 100 Hz from M1 ECoG signals in the 6-OHDA model animals (**Fig 1g**). We extracted the mean value of the frequency range 20-42 Hz, corresponding to the High Beta - Low Gamma band (HB-LG), which has been established as the pathological neural correlate in hemiparkisonian rats [27]. We modelled the group comparison using a Linear Mixed Model, setting the event type as the fixed effect and the day of recording nested within the subject variable as the random effect. We identified a distinct modulation of the HB-LG band across the locomotor states, highlighting an increased activity specifically during states of akinesia, compared to stationary movement (P *<* 0.001) and gait (P *<* 0.001) (**Fig 1h**).

These results confirmed that 6-OHDA rats exhibited prolonged period of akinesia compared to sham [28], and that HB-LG band activity correlates with motor symptoms [29]. The presence of the akinetic state in the PD rats, together with the increase in HB-LG band power during akinetic episodes, indicated the pathogenic characteristics of this state.

### 2.2 Linear deep phenotyping analysis expands neurokinematic features separating locomotor states

To expand our understanding of the neural correlates of locomotor deficits, we created a multimodal dataset for deep neurobehavioral phenotyping. We synchronized 36 kinematic features (from 3D coordinates recorded at 200 Hz) and 108 neural features sampled at 200 Hz (please refer to **Suppl. Table 1** for the full list of features used). Thus, the identical 5 ms interval could be examined from both a kinematic and a neural perspective, creating a vector of features that is continuous in time. We used *neurokin* to handle the extraction of kinematic features from 3D hindlimbs coordinates and synchronize neural features form ECoG data (extracted using *py_neuromodulation* [30]) (**Fig 2a**).

**Fig. 2.**
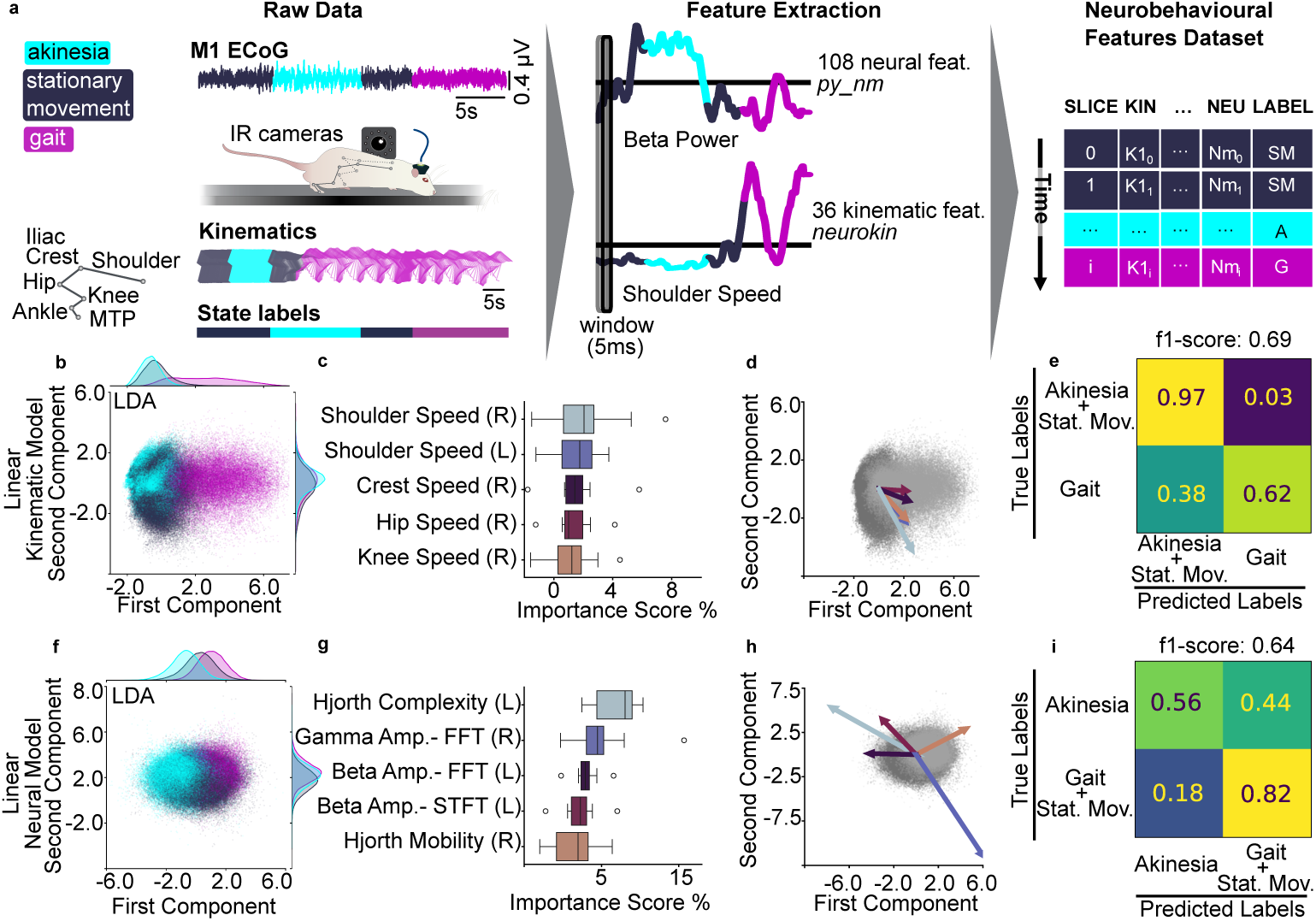
Linear deep phenotyping analysis expands neurokinematic features separating locomotor states. **a. Feature extraction.** Runway trials were synchronized across videos, 3D kinematics (IR cameras) and ECoG. We used *neurokin* to compute 36 kinematic features, and *py neuromodulation* to compute 108 neural features. Locomotor events were labelled offline. **b. LDA - kinematic model.** Gait clustered distinctly along LD1; akinesia overlapped stationary movement (marginal densities shown). **c. Top kinematic contributions.** Permutation test ranked speed variable highest. (body side indicated in brackets). **d. Kinematic loadings.** Higher speed-related variables correlated with the gait state. The length of the arrows is proportional to the weight on the components. **e. Confusion Matrix - kinematic model.** Cross-validated LDA separated gait from non-gait states. **f. LDA - neural model.** Akinesia shows modest segregation; gait and stationary movement largely overlap (marginal densities shown). **g. Top neural contributions.** HB-LG band power and Hjorth complexity led the ranking (hemisphere indicated in brackets). **h. Neural loadings.** HB-LG related metrics and Hjorth complexity biased toward akinesia; gamma power and Hjorth mobility toward gait. The length of the arrows is proportional to the weight on the components. **i. Confusion Matrix - neural model.** Cross-validated LDA discriminated akinesia above chance.

After extraction of the neurobehavioral feature vectors, we set to investigate specific features separating locomotor states. We created two linear feature spaces i) utilizing the kinematic portion ii) utilizing the neural portion of the multimodal dataset. In the kinematic space, the model could separate the gait state, but confounded the stationary movement with the akinetic state (**Fig 2b**). To understand which features contributed the most to the state separation, we inspected the model with a permutation feature importance test. We repeated the test in a cross-validation design, leading to the identification of the top 5 features (speed of: right shoulder, left shoulder, right crest, right hip and right knee) (**Fig 2c**). We visually inspected the directionality of these features by plotting the corresponding scaling on the LDA space. This confirmed that high values in speed-related variables intuitively correlated with the direction of separation of the gait state (**Fig 2d**), thus validating our pipeline. Finally, we quantified the performance of the model in predicting the state values of unseen data, within a cross-validation design. We then computed the f1-score (a metric combining precision and recall) which returned a value of 0.69, and calculated the average confusion matrix (**Fig 2e**). Because the kinematic model fails to separate akinesia from stationary movement, we grouped the two categories (**Suppl. Fig 1a** displays the expanded 3×3 confusion matrix). In line with the projection, the LDA model could separate akinesia and stationary movement (97% *±*3% true positive rate) from the gait state (62% *±*14% true positive rate).

Conversely, the LDA model based on neural data only, improved the separation of akinesia from stationary movement and gait, but confounded gait and stationary movement. (**Fig 2f**). We computed the most relevant neural features for states separation (**Fig 2g**), and identified Hjorth complexity and mobility [26] as relevant features, while also confirming modulation of HB-LG and gamma oscillations. Projecting the loadings in the LDA space (**Fig 2h**), suggested that Hjorth complexity in the left hemisphere (6-OHDA injected) correlated with akinetic episodes, while Hjorth mobility in the right hemisphere correlated with movement. Finally, we quantified the performance of the model in predicting the state of unseen data (f1-score:0.64), and computed the average confusion matrix (**Fig 2i**). Here, we grouped the categories of stationary movement and gait as our focus was the neural distinction between the akinetic state versus active states (**Suppl. Fig 1b** displays the expanded 3×3 confusion matrix). Indeed, the model returned a complementary result to the kinematic model: the akinetic state was distinguished above chance level (56% *±*11%) from the active states of gait and stationary movement (82% *±*6%).

### 2.3 Deep phenotyping with neural networks refines locomotor state characterizations

To account for possible limitations of linear models, such as LDA, for non-linear data [31], we additionally performed a non-linear analysis to find the neural features that best discriminate locomotor states. We used the same multimodal dataset, and the previously annotated locomotor state labels as ground truth. We employed the toolbox CEBRA [32], a deep neural network trained with contrastive learning. We computed the low-dimensional embeddings using supervised training for the selection of the contrastive learning sampling and then trained a K-Nearest Neighbors (KNN) classifier to estimate the performance of the model.

We first evaluated the kinematic portion of the features dataset. We projected the kinematic data in a low-dimensional embedding, to visualize the distribution (**Fig 3a**). Then, we assessed the importance of each feature within a cross-validation design. We embedded the test dataset, randomizing one feature at a time, and evaluated the embedding with a KNN classifier by computing the f1-score. Calculating the f1 importance score allowed us to estimate what kinematic features are most relevant in constructing the embedding (**Fig 3b**). As for the linear analysis, speed was the predominant feature category. We mapped the value of kinematic features onto the data points, creating features heatmaps. to inspect how the features mapped into the embedding (**Fig 3c**, and **Suppl. Fig 3a**). Finally, we evaluated the performance of the model in classifying unseen data by computing the average confusion matrix (f1-score: 0.87), based on a KNN classifier (**Fig 3d** and **Suppl. Fig 2a**). The classification score maintained a good level of separation (94% *±*8% true positive for the akinetic/stationary group, 83% *±*13% true positive for the gait class), greatly improving on the false negative score for the gait state (17% *±*13% false negative), at the expense of a small rise in its false positive score (6% *±*8% false positive) in comparison to the LDA model (**Fig 3d**),

**Fig. 3.**
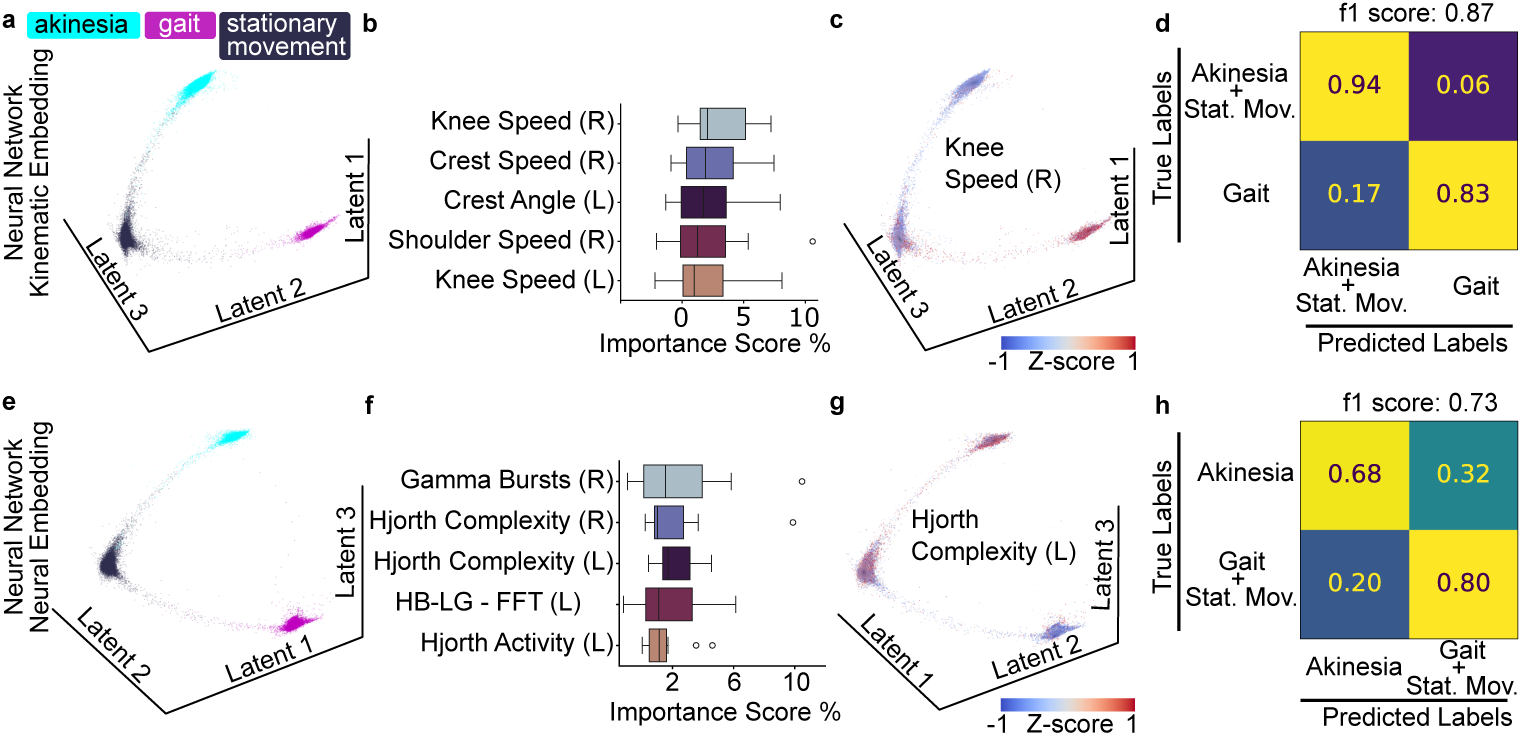
Deep phenotyping with neural networks refines locomotor state characterizations. **a. CEBRA embedding - kinematic model.** Supervised CEBRA embedding of 36 kinematic features separated locomotor states. **b. Top kinematic contributions.** Permutation test on a KNN classifier ranked speed-related variables highest **c. Kinematic feature projection.** Feature mapping in the embedding space showed high right knee speed clustering in gait points **d. Confusion Matrix - kinematic model.** KNN on the embedding achieved precise gait separation **e. CEBRA embedding - neural model.** Supervised CEBRA embedding of 108 neural features separated locomotor states. **f. Top neural contributions.** Permutation test on a KNN classifier ranked gamma bursts and Hjorth parameters and HB-LG amplitude variables highest **g. Neural feature projection.** Elevated Hjorth Complexity mapped onto akinesia. **h. Confusion Matrix - neural model.** KNN on the embedding achieved precise akinesia separation.

Next, we performed the same non-linear analysis, using only the neural portion of the multimodal dataset. After supervised training, we transformed the data onto a 3D embedding (**Fig 3e**), which visually indicated a better separation of the states’ clusters than achieved by the linear analysis with LDA (**Fig 2f**). Computing the f1 importance score validated the relevant role played by Hjorth complexity, which was previously suggested by the linear model (**Fig 3f**) and of HB-LG and gamma amplitude. Mapping the values of each parameter onto the data points revealed a cluster of high Hjorth complexity values, in correspondence of the akinetic state, and partially the stationary movement state (**Fig 3g**, and **Suppl. Fig 3b**). Hjorth complexity showed a similar embedding mapping as the HB-LG band amplitude feature (clustering over akinetic states); in contrast, the gamma power maps with gait and partially stationary movement. Finally, computing the confusion matrix of the classification of unseen data (f1-score:0.73), revealed an improvement in the classification of akinetic states (68% *±*9% of true positive for akinetic class and 80% *±*7% for the gait/stationary movement group) an improvement in the false negative (32% *±*9%), at the expense of a slight rise in the false positive score (20% *±*7%) in comparison to the LDA model (**Fig 3h** and **Suppl. Fig 2b**).

### 2.4 Identified neural features modulate during akinetic state

Next, we set out to characterize the identified neural features relevant for the separation of locomotor states, and their relationship to akinetic episodes. Plotting each feature as a function of its importance score for the linear and deep-learning models, allowed us to identify consistently high-scoring features: Hjorth complexity (left hemisphere), gamma amplitude (right hemisphere), Hjorth mobility (right hemisphere) and HB-LG amplitude (left hemisphere) (**Fig 4a**). Hjorth complexity and mobility reflect, respectively, the deviation from a pure sine wave, and the average frequency content [26] as graphically represented in **Fig 4b**. Plotting the average temporal modulation of these four features around the onset of (N=75) episodes of akinesia revealed common patterns. Specifically, Hjorth complexity underwent a gradual increase shortly before the beginning of the akinetic episode, and again at the onset (**Fig 4c**). Conversely, Hjorth mobility displayed a gradual decline preceding and during the akinetic episode onset (**Fig 4d**). The HB-LG band showed a similar modulation profile as Hjorth complexity in the affected hemisphere, whereas gamma showed a similar modulation as Hjorth mobility in the healthy hemisphere. We then explored the group-level modulation of these features across locomotor states. We calculated significance with a related t-test (*α <* 0.05), adjusting for multiple comparisons using a Bonferroni-Holm correction (12 tests). Hjorth Complexity was significantly up-modulated specifically during states of akinesia, whereas gamma amplitude demonstrated a down-modulation during episodes of akinesia, in comparison to the gait state (**Fig 4e**). Finally, we qualitatively inspected the ability of the HB-LG amplitude or Hjorth complexity to segregate the locomotor states. We computed two embeddings based only on one of the two neural features. Both the data projection in the embedding and the f1-scores (calculated in a cross-validation design) show comparable separation, with a slight advantage of the Hjorth complexity over the HB-LG band-based embedding (**Fig 4f**). Together, our results highlight Hjorth complexity and mobility as possible real-time markers for state and state transition detection.

**Fig. 4.**
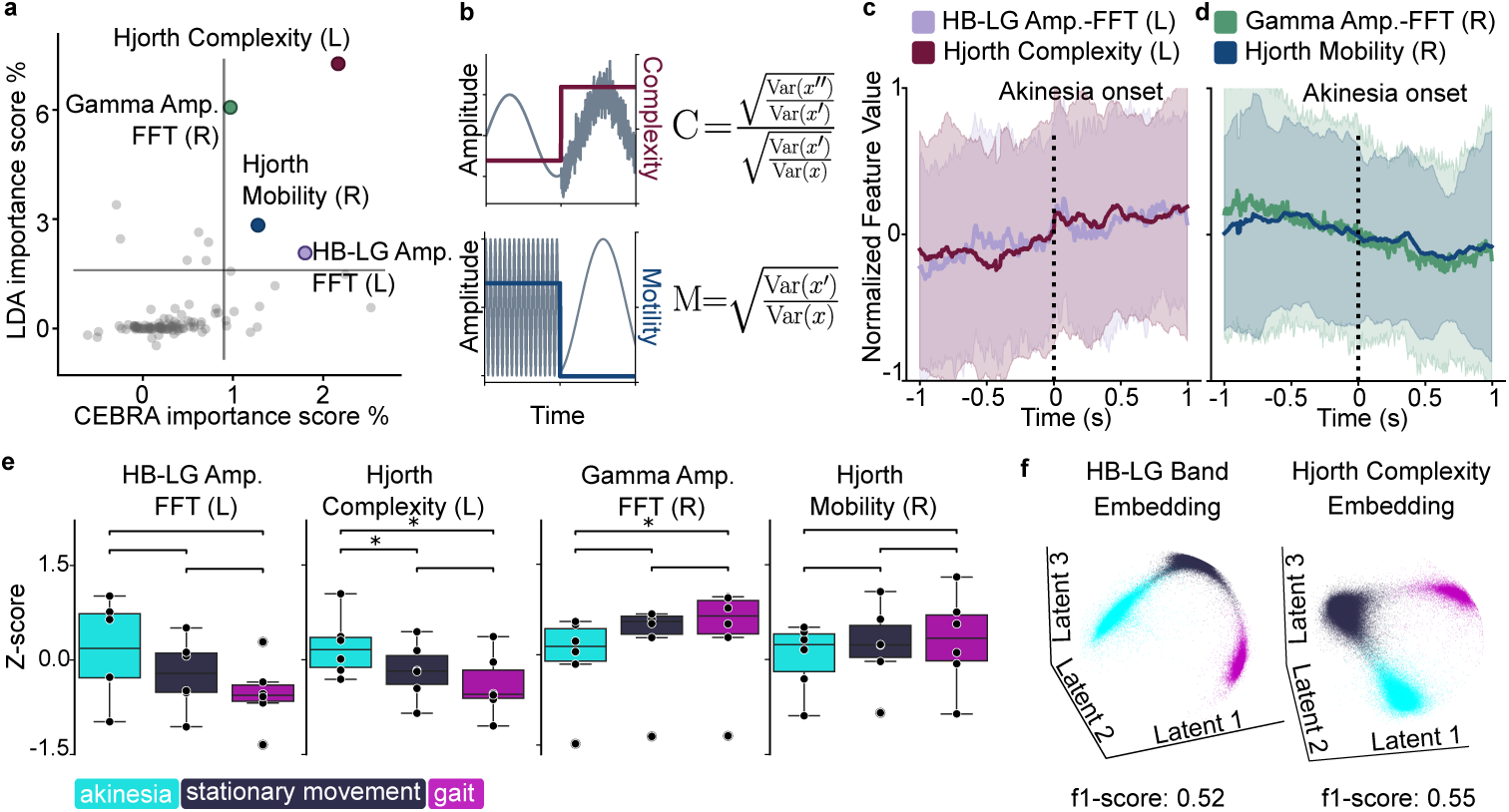
Identified neural features associate with onset of akinetic state. **a. Joint features ranking.** Scatter of LDA vs CEBRA importance highlighted shared top-scoring features. **b. Hjorth parameters schematic.** Visual example of modulation in the Hjorth parameters **c. Hjorth complexity positively correlated with akinetic onset.** Mean ± SD curves showed Hjorth complexity and HB-LG power increasing on akinesia onset. **d. Hjorth mobility negatively correlates with akinetic onset.** Mean ± SD curves showed Hjorth complexity and gamma power decreasing on akinesia onset. **e. Group effects.** Akinesia correlated with increased Hjorth complexity and decreased gamma amplitude (paired t-test, Holm-corrected for 12 tests). **f. Single feature embeddings.** Left: CEBRA embedding created using only the HB-LG Amplitude - FFT (L) feature. Right: CEBRA embedding created using only the Hjorth Complexity (L) feature.

### 2.5 Modulation of identified neural correlates is confirmed during freezing of gait in a Parkinson’s patient

Finally, we investigated whether the identified features (Hjorth complexity and mobility, beta (13-20 Hz) and gamma amplitude) could translate to the characterization of locomotor deficits in Parkinson’s patients. For this, we analysed kinematic and bilateral STN data synchronized with manual gait and freezing of gait (FoG) labels from video recordings. The patients were instructed to walk in a straight line for 10 meters, turn and walk back (**Fig 5a**). An exemplary episode of FoG is depicted in (**Fig 5b**). We binned the data in 1 second bins, and selected lower-body markers: lateral condyles, shank, V. metatarsal bone, toe. Using *neurokin*, we computed a set of 292 kinematic features from the following categories: linear velocity, linear speed, linear acceleration, tangential acceleration, angle, angle velocity, angle acceleration, toe height, toe forward movement. Next, we performed a PCA to quantitatively assess the distinct kinematic features of FoG to normal gait (**Fig 5c**). Picking the top 10 correlated features returned a selection of minimum acceleration features which negatively correlated with gait, and maximum velocity or speed which positively correlated with the gait state(**Fig 5d**). We then analysed the STN neural data, segmenting episodes of gait and FoG. To asses clinical translation of the neural features identified in the 6-OHDA model, we computed the amplitude of Hjorth complexity, Hjorth mobility, beta band, and gamma band. We analyzed a total of 6 STN channels (3 left hemisphere, 3 right hemisphere) and for each participant displayed an exemplary channel, which showed the maximum beta modulation during FoG. While one subject did not display any neural modulation, the second subject showed the same pattern of modulation of Hjorth complexity and mobility during FoG, as observed for M1 ECoG during akinesia in our rodent model (**Fig 5e** and **Suppl. Fig 4**). Specifically, Hjorth complexity was up-modulated (P=0.004), Hjorth mobility was downregulated (P=0.004) the beta band was up-modulated (P=0.006), following the same modulation as for 6-OHDA rats. However the gamma band was up-modulated (P=0.004). We used a within-patient, trial-constrained permutation test, with Bonferroni correction for 4 tests. These observations establish the capability of deep neurobehavioral phenotyping to identify features for locomotor states separation that translate in clinical experimental settings.

**Fig. 5.**
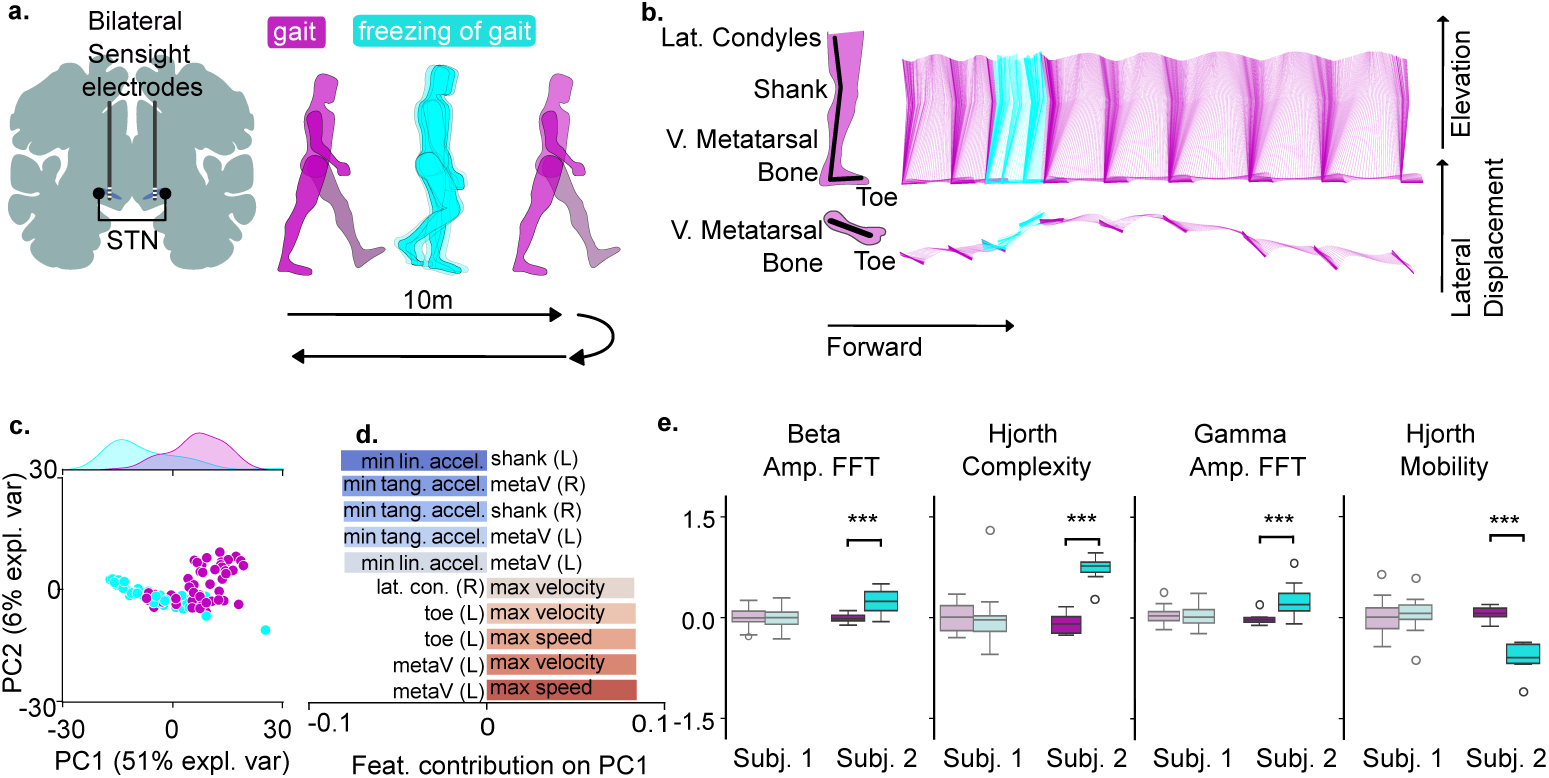
Newly identified neural correlates of akinesia confirm modulation in a Parkinson’s patient Experimental set up. Patients performed a 10 meters walking test. Kinematic and neural data (STN) were synchronously collected during the experiment. **b. Kinematic diagram of exemplary FoG episode** Selected body parts for kinematic analysis (left), stick diagrams (right) **c.**and **d.** PCA reveals main modulated kinematic features during FoG **e. Neural correlates of FoG** Statistical analysis showed significant correlations of Hjorth complexity and mobility in one patient for FoG episodes.

## 3 Methods

### 3.1 *neurokin*’s modular and open-source design

To conduct tasks of synchronization, data parsing and kinematic feature extraction, we made use of the self-developed package *neurokin* which streamlines the integration of three distinct types of data: neural local field potentials, kinematic recordings, and event labels. The package is primarily object-oriented, to ensure a coherent interface across data types. The underlying principle of *neurokin* is modularity, both enabling the use of any of the classes independently (to facilitate data import and management), and supporting the assembling of project-specific pipelines.

We sought to adhere to the FAIR (Findable, Accessible, Interoperable, Reusable) principles [33]. Specifically, *neurokin* is openly available on GitHub ([neurokin](https://github.com/ELGarulli/neurokin)) and pip installable ([pip install neurokin](https://pypi.org/project/neurokin/)) (”**F**” principle), it includes independent documentation ([neurokin documentation](https://neurokin.readthedocs.io/en/latest/)) (”**A**” principle), it uses standard domain-relevant file types (e.g., .c3d or .csv) (”**I**” principle), it builds upon qualified software (e.g., numpy, pandas) and details a permissive MIT License (”**R**” principle). Building on these principles, we encourage community-wide code review and participation for further development.

#### KinematicDataRun

This class was used for importing and processing kinematic data. It handles 2D and 3D kinematic data originating from optoelectrical gait systems such as Vicon (.c3d file format), or files from computer vision toolboxes (e.g., DeepLabCut). Upon importing the data in the corresponding kinematic object, the user can make use of the kinematic-features extraction pipeline. This pipeline was designed to maximize flexibility. To this end, the user can define a custom skeleton via a configuration file, and select standard features (e.g., speed, acceleration, height). Additionally, the framework supports the straightforward addition of custom features.

#### NeuralData

This class operates on neural recordings, such as ECoG and Local Field Potentials (LFPs), standardising data from different recording systems in a common format to interface with fundamental methods (e.g., timestamps-parsing, PSD calculation). It is currently compatible with the Tucker-Davis Technologies apparatus and OpenEphys data, with the possibility to be expanded to other hardware. It was used for importing and processing of the collected neural data from rats

#### NeuralCorrelatesStates

We used this class to tailor a pipeline to retrieve Vicon labels and align them with kinematic and neural features. This can easily be expanded to support any time-stamped, user-defined events. Users can define the events by providing labels. These are used to retrieve corresponding kinematic and neural data, and can eventually be used as inputs for machine learning applications.

### 3.2 Animal Model and Surgical Procedures

All experiments were conducted in accordance with the German regulatory body (LaGeSo), specifically under the animal licence number G0206/19. Experiments were performed on male Sprague Dawley rats with initial weights of 200-250 g, ordered from Janvier Labs. Animals were kept in a 12-hour light cycle with ad-libitum access to food and water.

All the interventions were performed under full general anaesthesia with isoflurane (1–2%) in oxygen-enriched air. After surgery, the rats were placed in an incubator for optimised recovery from anaesthesia as described in [34].

PD models were injected with 6-OHDA (Sigma: H4381) dissolved immediately before use in a buffer solution of 0.02% w/v ascorbic acid achieving a final concentration of 8 mg/mL. Of the 6-OHDA solution 1.2 *µ*L was injected at a rate of 0.04 mL/min through a pipette according to [16]. Shams were injected with the same solution except for 6-OHDA. The injection was targeted to the Medial Forebrain Bundle (coordinates from bregma: AP: −2.6 mm, ML: 1.6 mm, DV: −8.4 mm from skull surface). The success of the 6-OHDA model was verified with the antibodies anti-tyrosine hydroxylase (abcam: ab112) and anti-rabbit Alexa 488 (Thermofisher: A-11034).

Following a recovery period of 2 weeks, the rats were implanted with 2 stainless steel ECoG screws (2 mm) over the motor cortex (left and right hemisphere), one ground screw over the cerebellum, and 2 LFP probes targeted to the STN area (left and right hemisphere). No data from the LFP electrodes was analyzed in this study.

To guarantee the protection of the implant, we designed a three component cap, that can be 3D printed and assembled during surgery. The 3D-printing files are openly available ([headstage caps](https://github.com/ELGarulli/headstage_cap.git)).

### 3.3 Runway Task

We trained the rats to run from one end of a 15×140 cm runway to the other. We placed a cage with at least one additional rat at the end of the runway to further motivate the tested rat. At the beginning of each recording the rat was placed at the starting end of the runway, then given up to a minute to spontaneously initiate walking. If, by the end of the given time, the rat had not run the whole length, it was gently brought to the ending side for a few seconds while the recording setup was prepared for the next recording. This ensured exposure to the additional rats and minimized idle time at the starting point. We repeated the task until 15 successful crossings were achieved, or the total recording time per animal reached 20 minutes. To ensure consistency across sessions, we marked with a permanent marker the joint points (shoulder, iliac crest, hip, knee, ankle, and metatarsophalangeal joint) on the skin. Then, at the start of each session, 4 mm reflective markers were glued with double-sided tape on the marked spots.

### 3.4 Kinematic Data Recording

To capture the movements of the rats we used an infrared camera-based motion capture system (Vicon). We equipped a 2.5 m2 room with 10 infrared and 2 HD wallmounted cameras. The infrared cameras were positioned around 3 sides of the runway covering different heights. Data from the infrared cameras was recorded at a sampling rate of 200 Hz, and videos from the HD cameras were recorded at 50 Hz. The raw data was then processed using the Nexus software (v2.12.1). Each recording was cropped from the earliest point where all the markers were visible and no contact between the experimenter and the animal was observed, to the last point where the markers were visible and no contact with the experimenter was observed, or to the end of the run. The 3D position of each joint was manually reconstructed using a custom-defined bipedal skeleton (as no markers were placed on the forepaws). The data was then exported in the .c3d format.

### 3.5 Neural Data Recording

To record ECoG neural signals we used stainless steel screws (DIN 84 A2 M1.2×2 from MaxWitte GmbH), laser soldered to a 10T stainless steel PFA-coated wire (SS-10T/A from Science Products). The wires, soldered to Omnetics 32-channel connectors, were coated with Epoxy resin to grant stability, and all the unused channels were shorted to the ground. During experiments, we plugged a digitising headstage (Intan), to minimise movement artefact and used a TDT recording apparatus. The signal pipeline was then controlled through the software Synapse. We recorded all raw data at a sampling frequency of 24414.1 Hz, data from implanted LFP electrodes were not analysed.

### 3.6 Locomotor States Labelling

We defined akinesia on a phenotypic level, observing an almost complete lack of any type of movement. This was to be contrasted with stationary movement, where the rat was in an engaged state and performing an array of behaviors (e.g., grooming or sniffing) and gait, when the rat was walking. The events, were manually labelled in the Nexus software. The data was then exported in the ASCII format to a .csv file.

### 3.7 Locomotor States Distribution Analysis

We used *neurokin* to process the .csv files generated by the Nexus software, and created a dataset summarizing all the events timestamps in each run. For each run, we computed the percentage of time spent in each locomotor state. Next, we computed subject-level averages and quantified the differences in locomotor state distribution between the sham control and the PD animals. We performed an independent t-test with a Bonferroni-Holm correction for multiple comparisons (3 comparisons). We assumed an *α <* 0.05.

### 3.8 Kinematic Features Dataset Creation

To create the neural and kinematic dataset we relied on a consistent folder structure that was then automatically processed. We used *neurokin* to import and process the .c3d and .csv files generated by Vicon. We selected kinematic features representing speed, acceleration, angle variation for the various markers, forward movement and height for the metatarsophalangeal joint. We based our feature extraction on a sliding-window strategy. The kinematic features were computed at a 200 Hz rate (i.e., 5 ms window with a 2.5 ms overlap with the following window).

### 3.9 Neural Features Dataset Creation

Using *neurokin*, we selected the neural data corresponding to the valid portion of the recording (all markers visible, no contact with the experimenter). To extract neural features we relied on the package *py neuromodulation* [30], and used a feature sampling rate of 200 Hz, thus aligning with the kinematic features framerate. We computed 108 parameters, among which temporal features (e.g., Hjorth parameters), oscillatory features (e.g., power of the beta band and gamma band) and waveform shapes (e.g., sharp wave intervals). In rare occasions, we encountered a length mismatch between the kinematic and the neural files. Upon reviewing the original files, we discovered that seldomly a few additional frames were recorded at the beginning of the kinematic file, we discarded files with excessive length mismatch, and trimmed those with minimal mismatch (typically 1-2 frames).

### 3.10 Multimodal Dataset Creation and Testing Parameters

For each of the valid runs, we filtered for outliers (set to median value) and Z-scored column-wise. Finally, we concatenated all runs in a multimodal (kinematics, neural and labels) dataset. To validate the model, we performed an 11-fold cross-validation - as 55 runs were present, at each iteration we trained on 50 runs and tested on 5 unseen ones. We used random seeds to ensure all analyses were done on the same splits.

### 3.11 LDA analysis

The visualization of the entire model was achieved by training and transforming the whole dataset and computing kernel density estimation marginal distributions. Importance scores were computed for each feature, for each fold of the cross-validation, using the *sklearn* package [35], specifically calculating the balanced accuracy. The final confusion matrix was calculated as the average of the 11 folds.

### 3.12 CEBRA analysis

We used a CEBRA model [32] with the following parameters: *model architecture* = “offset200-model”, *batch size* = 512, *temperature* = 0.1 (value set after temperature optimization scan), *learning rate* = 0.0005, *max iterations* = 50000, *time offsets* = 200, *output dimension* = 3, *conditional* = “time delta”. This allowed us to use the state labels as the categorical variable used to optimize the contrastive learning steps. The “offset200-model” was custom made to have a receptive field of 200 samples, corresponding to 1s of data. To evaluate the performance of the model we used a KNN decoder with *n neighbors* = 3 and *metric* = “cosine”. Importance scores were computed for each feature, for each fold of the cross-validation, using the *f1 score* calculation from the *sklearn* package. The final confusion matrix was calculated as the average of the 11 folds.

### 3.13 Statistical Analysis of Selected Features Alignment

The significance of each selected feature, was computed as a related t-test between the compared state. To correct for multiple comparisons, we used a Bonferroni-Holm correction, ranking 12 t-test (3 states, 4 features). We assumed an *α <* 0.05.

### 3.14 Human data collection

We analyzed the data of two male parkinsonian patients (Age: 68 and 70 years) implanted bilaterally in the Subthalamic Nucleus (STN-DBS) with Sensight B33005 electrodes and the Percept PC implantable pulse generators (Medtronic, PLC, USA). Patients showed gait disturbances and freezing of gait in meds-off/stim-off condition (i.e., after overnight suspension of all dopaminergic drugs and after switching off the stimulation for at least one hour), remarkably improved by both medication and stimulation (Unified Parkinson’s Disease Rating Scale, UPDRS-III [36]), gait item: i) meds-off/stim-off: 50 and 78, ii) meds-off/stim-on: 31 and 45.

Data were collected in the context of the project 24778381 within the Collaborative Research Centre TRR 295 ReTune (Deutsche Forschungsgemeinschaft [DFG]), which received ethical approval from the institutional review board at Würzburg University (n. 103/20-am) on 22.03.2021. The project conformed to the declaration of Helsinki and written informed consent was obtained from the participants at recruitment.

Patients performed the experiment in meds-off/stim-off condition. They were instructed to walk barefoot back and forth for 10 times along a walkway of about 10 meters, at their preferred speed. Kinematic data were recorded using a six-camera optoelectronic system (SMART DX-400, BTS Bioengineering, Italy) with a sampling rate of 100 Hz and 29 markers placed on anatomical landmarks according to a published full-body protocol. In-built visible-light camera (VIXTA, BTS Bioengineering, Italy) allowed to have automatically synchronized videos of the experiment. LFP were collected bilaterally and synchronously from all non-adjacent contact pairs (i.e., [0–2], [1–3], [0–3], being the 0 the lowermost and 3 the uppermost contact, recorded with the indefinite streaming modality) with a sampling frequency of 250 Hz. Cortical activity was recorded with a 64-channel portable EEG device (Sessantaquattro, OTBioelettronica, Italy) here used just for synchronization purposes. The kinematic and cortical recordings were synchronized using a transistor-transistor logic (TTL) reference signal, while cortical and LFP recordings were synchronized by means of an electrical artefact produced by a transcutaneous electrical nerve stimulation (TENS), as described in detail elsewhere. In this way, we could align kinematic data and labels from videos with the LFP time series.

Videos were inspected independently by two clinicians with experience in movement disorder. FOG episodes were recognized when patients either failed to start walking within one second after being instructed to do so, or showed visible signs of attempting, but being unable, to continue walking. Additionally, any sudden and noticeable interruption in walking without an obvious external cause was also classified as a FOG event [4]. Beginning and end times of each FOG episode was marked independently, and any discrepancies between the two experienced clinicians were resolved through mutual discussion. Subject 1 had 20 freezing episodes, during turning and continuous walking (median duration: 7 seconds, range 1-33 seconds). Subject 2 had 8 freezing episodes mainly during turning (median duration: 3.5 seconds, range 2-8 seconds).

### 3.15 Human data analysis

We processed the 3D kinematic data by parsing the moments of straight walk (as the markers were not detected during the turns) and flipped the direction of the backwards walks to be able to compute equal features. We selected markers that had at least 70% of the points and interpolated the missing values. We chose a subset of markers in the lower body and computed a set of 292 linear and angular features using *neurokin*. We filtered for outliers (set median value) and Z-scored column-wise. We split each run in portions of 1 second and computed the mean for each portion. We performed a dimensionality reduction step using PCA and finally color-coded depending on the freezing of gait label (if the portion of data spanned a transition, the color was decided by the majority of data).

We processed the STN data by selecting the valid part of the data (exclusion of the flanking zero-padding) and computed neural features using *py neuromodulation* resampled at 100 Hz. For visualization purposes we selected the channel which had the highest beta band modulation and was coherent with the trend (see **Suppl. Fig 4** for the complete set). We excluded a short segment of the data (6 seconds)around a strong STN artefact from session 2 of Subject 2. We parsed the data in episodes of FoG and of gait (Subj. 1 - gait episodes: 20, FOG episodes: 19; Subj. 2 - gait episodes: 9, FOG episodes: 8). Finally, the mean of each feature per episode was computed.

## 4 Discussion

In this study we set out to discover novel neural fingerprints of akinetic states, by analysing kinematic and ECoG multimodal data from a Parkinson’s rodent model. We created a neurobehavioral dataset, capturing both the kinematic and neural aspects of locomotor states. To identify critical neural features characterizing akinetic states, we trained a linear model and a neural network on the neural and kinematic portions of the dataset. Specifically, our results linked Hjorth complexity and Hjorth mobility to akinetic episodes. Further, we confirmed the same pattern of modulation in one of two Parkinson’s patient, thus supporting the potential for translational application. We propose these novel neural fingerprints as potential biomarkers of akinetic states, which could be further explored as targets for precision interventions such as closed-loop stimulation.

### 4.1 Optimized feature analysis for akinesia characterization

Designing algorithms to detect FoG is an on-going effort, even inspiring international Machine Learning challenges [37]. Previous literature focused on classifying FoG computing features based on the gait cycles [38–40], which is well-suited for classifying gait abnormalities but entails intrinsic delays, as detection can only occur after the completion of a step. Moreover, gait-cycle-based approaches cannot easily distinguish akinesia from voluntary resting states, as both lack stepping events. To overcome these limitations, we employed a rolling time-window approach [41, 42], in line with recently developed motion sequencing algorithms [43]. This enabled continuous monitoring of locomotor states, including completely akinetic episodes. Moreover, leveraging a neurobehavioral dataset, presented two different lenses (neural and kinematic), on the same phenomena.

### 4.2 Data-driven discovery of translatable features

Parkinson’s disease is characterized by a heterogeneous symptom distribution [44–46]. Historically, the most explored feature has been the beta band, which has shown robust correlation with bradykinesia, rigidity and tremor [13–15]. However, a single feature might not account for the complexity of different symptoms distribution [47]. To address this, we computed a broad set of neural and kinematic features and employed a data-driven approach to select emerging salient features characterizing pathological locomotor states. We retrieved known correlates of akinesia and bradykinesia such as beta band amplitude [48, 49], as well as known correlates of prokinetic activity such as gamma band amplitude [50], thereby corroborating the validity of our method. Interestingly, less explored but promising [51] features such as Hjorth complexity and mobility, were also significantly modulated. To probe the translational value of our findings, we also explored a small human dataset. While one subject did not show any pattern of modulation (neither in the Hjorth parameters, nor in the hallmark beta band), the second highlighted a modulation pattern consistent with our findings. Together, these results support our data-driven pipeline as an agnostic approach for identifying salient neural features, revealing both established and novel markers of pathological locomotor states across species.

### 4.3 Hjorth Parameters as Novel Time-Domain Biomarkers of Akinesia

Hjorth parameters were first established to quantify EEG traces, and they provide a bridge between the time domain (the signal change over time) and the conventionally analysed frequency domain (the signal composition over frequencies) [26]. Since then, they have been explored as biomarkers to classify Alzheimer’s Disease patients [52, 53], classify Parkinson’s Disease patients [54] and to predict tremor in Parkinson’s patients [55]. Here, we established that Hjorth parameters correlated with a pathological locomotor state, and uncovered the specific modulation of the complexity and mobility parameters at the onset of akinesia, both in a PD rodent model and in a patient. Time-domain biomarkers like Hjorth parameters are appealing for realtime closed-loop stimulation because they bypass the added computational cost of frequency-domain analyses and the latency inherent to causal filters [14].

### 4.4 Translational comparability of gait interruptions in rodents and human

Freezing of gait (FoG) in Parkinson’s disease remains a poorly understood phenomenon [56, 57]. Whether the proposed mechanisms of FoG in human translate to rodent models requires consideration at multiple levels, including neuroanatomy, motor behavior and network activity. Several circuit pathologies are hypothesized to play a role in the manifestation of FoG, including distortions in locomotor networks (i.e. cortex-basal ganglia loops and brainstem MLR), their cognitive regulation (i.e. frontal cortex and visual cortex), and their emotional control (i.e. amygdala) [56–60]. Rodent models of PD typically rely on dopaminergic depletions in the nigrostriatal pathway [22]. The resulting circuit deficits in rodents do reflect the progressive dopaminergic degeneration in patients that correlate with the occurrence of FoG [61]. The neuroanatomical comparability of our study is therefore established predominantly at the level of dopaminergic deficiencies acting on cortex-basal ganglia loops. At the behavioral level, there are four major motor expressions of FoG in patients, namely: leg trembling, shuffling, festination and akinesia [3]. In our rodent experiments, we only observed akinesia as a distortion to the natural gait sequence on the runway. These observations are consistent with an in-depth literature review that found no clear evidence for gait interruption patterns in rodent models of PD other than akinesia [22]. There has been one report in the non-human primate MPTP model, where leg trembling occurred in the hindlimbs during quadrupedal locomotion [62]. This observation indicates that other movement patterns - resembling human FoG - can occur during quadrupedal gait. As a conservative interpretation of our study, we conclude that rodent models of PD and human patients are comparable in the sense that they spend excessive amounts of time in pathological motor states during gait execution (i.e. akinesia in rodents and multiple patterns of FoG in patients). Within these pathological motor states, we observed several similarities in the network oscillations of cortex in rodents and STN in one human patient. Namely, beta oscillations, as well as Hjorth complexity and mobility modulated the same direction in rodents and one Parkinson patients during pathological gait states. For beta oscillations, these findings have already been substantiated by fitting clinical evidence in a group of 18 Parkinson patients [63]. Yet, future research will need to address the reproducibility of the modulation of Hjorth parameter modulations during gait in larger patient cohorts. Establishing translationally robust neural biomarkers will be pivotal when developing next generation neuromodulation technologies.

### 4.5 Towards multi-input closed-loop stimulation control

The ideal therapy should be tailored to the specific subset of symptoms that a patient displays, maximizing the benefits while minimizing adverse effects. Thus, finding a range of computationally lightweight features, that can be combined in a multi-input closed-loop control [21] and can reliably detect a pathological state will be crucial to develop personalized stimulation strategies to mitigate, or avoid altogether a pathological motor state. A feature vector that allows to identify a pathological state, can be conceived as discovering its fingerprint, which could eventually be targeted with a timely neuromodulation. Recent examples of model-based control are offered in the field of Spinal Cord Injury rehabilitation, where a context aware algorithm could significantly improve locomotor performance [34, 64]. The practical need of a state-based stimulation is illustrated, for example, by the impairment of movement stopping caused by a state-naive open-loop STN stimulation to address Parkinson’s symptoms [65]. Ultimately, we envision a closed-loop algorithm based on real-time mapping of the neural state projected to a patient-specific embedding. This tailored neural projection would reveal the transitions into pathological states and guide targeted interventions to relieve or avoid pathological manifestations. As a proof of concept, we tested the monitoring abilities of our neural embedding in an offline implementation. **Video supplement 2** illustrates a successful neural state tracking during a runway trial.

## 5 Limitations

The analysis presented in this study was focused on three locomotor behaviors (akinesia, stationary movement, and gait) that were clearly recognizable during the runway task. It is notable that both the linear and non-linear kinematic models could not perfectly separate the akinetic state from the stationary movement. However, this is explainable as no markers were placed on the forelimb or on the head of the animals, which often remained the only body parts active during stationary movement. So, while the akinetic state was apparent from the video recording (used to create the ground-truth labels), it was challenging to distinguish from the available 3D coordinates. Further analyses of this Parkinson’s model could include other behavioral states, and potentially lead to the uncovering of additional neural correlates of the disease. Nevertheless, the pipeline established here could serve as a template to explore other states, as well as the effects of treatments (e.g., pharmaceuticals, electro- or optostimulation) on neural correlates.

The modulation of Hjorth parameters was reproduced in a single patient, although consistent and promising, more data needs to be collected.

## 6 Outlook

In conclusion, this study employed a data-driven approach to uncover unexplored neural correlates of pathological locomotor states in Parkinson’s disease across species. In the future, these biomarkers may prove to be novel targets for a closed-loop stimulation that adapts in real-time to the current locomotor state, providing an on-demand delivery method.

## Supplementary information

We have included an additional file containing the supplementary figures and tables mentioned in the manuscript. Additionally, we have submitted two supplementary videos.

## Acknowledgements

Computation has been performed on the HPC for Research/Clinic cluster of the Berlin Institute of Health. We thank Gianni Garulli for help in developing the 3D-printable recording cap. We thank the Scientific Laboratories Center at Charité for 3D-printing services. We thank Dipl.-Ing. Raik Paulat for technical support and set-up of the recording room. ELG, TM, BK, DD, CP, WJN, YP, ME, CH and NK received funding from Collaborative Research Center ReTune TRR 295-424778381. WJN received funding from the European Union (ERC, ReinforceBG, project 101077060), and the Bundesministerium für Bildung und Forschung (BMBF, project FKZ01GQ1802). M.E. received funding from DFG under Germanýs Excellence Strategy – EXC-2049 – 390688087, Clinical Research Group (KFO) 5023 BeCAUSE-Y, project 2 EN343/16-1. BMBF, DZNE, DZHK, DZPG, EU, Corona Foundation, and Fondation Leducq. C.H. received funding from DFG under Germanýs Excellence Strategy – EXC-2049 – 390688087 (SPARK-BIH Stroke Protect), EraNet Neuron project DFG project number 522473931, DZHK/BMBF Innovation Cluster (FKZ81X2100285), Charité 3R (C3RHeaD and STROKE-PREDICT-C3R), BIH, and Fondation Leducq Transatlantic Network of Excellence (17CDV03).

## Declarations

W.J.N. received honoraria for consulting from InBrain – Neuroelectronics that is a neurotechnology company and honoraria for talks from Medtronic that is a manufacturer of deep brain stimulation devices unrelated to this manuscript.

## Ethics approval

All experiments were conducted in accordance with the German regulatory body (LaGeSo) under the animal licence number G0206/19.

## Data availability

The full dataset analysed in this manuscript is published with the DOI: **10.5281/zenodo.15163493**. 3D models of the recording cap are available on GitHub ([headstage caps](https://github.com/ELGarulli/headstage_cap.git))

## Code availability

The full code used to perform all the analysis in this manuscript is available in the [akinesia fingerprint article repository](https://github.com/ELGarulli/akinesia_fingerprint_article). The *neurokin* toolbox is available on GitHub ([neurokin](https://github.com/ELGarulli/neurokin)) and pip installable ([pip install neurokin](https://pypi.org/project/neurokin/)).

## Author contribution

E.L.G., and NW conceived of the presented idea. E.L.G, T.M. and Y.P. gave critical input and guided the analysis, E.L.G. performed the analysis. E.L.G prepared figures and graphical illustrations. E.L.G, G.E.H., D.D. and M.S. developed neurokin. E.L.G, B.K. and R.B. annotated the videos and reconstructed the 3D coordinates. E.L.G, B.K., and R.B. carried out the experiments. B.K., R.B., A.V. and N.W. established the 6-OHDA Parkinson’s model. B.K., R.B. and A.V. performed TH staining experiments. B.K., R.D.S. and N.W. performed the surgeries. CP and IH collected and preprocessed the human data. E.L.G wrote the manuscript with support from N.W.. T.M., D.D., M.S., I.H., C.P., W.J.N., Y.P. M.E., C.H. and N.W. gave critical comments to the final manuscript. M.E., C.H. and N.W. supervised the project.

## References

[1] Martínez-Fernńndez, R., Schmitt, E., Martinez-Martin, P. & Krack, P. The hidden sister of motor fluctuations in parkinson’s disease: A review on nonmotor fluctuations. Movement Disorders 31, 1080–1094 (2016). URL 10.1002/mds.26731.

[2] Weintraub, D. & Burn, D. J. Parkinson’s disease: The quintessential neuropsychiatric disorder. Movement Disorders 26, 1022–1031 (2011). URL 10.1002/mds.23664.

[3] Nutt, J. G. et al. Freezing of gait: moving forward on a mysterious clinical phenomenon. The Lancet Neurology 10, 734–744 (2011). URL 10.1016/S1474-4422(11)70143-0.

[4] Schaafsma, J. D. et al. Characterization of freezing of gait subtypes and the response of each to levodopa in parkinson’s disease. European Journal of Neurology 10, 391–398 (2003). URL 10.1046/j.1468-1331.2003.00611.x.

[5] Bloem, B. R., Hausdorff, J. M., Visser, J. E. & Giladi, N. Falls and freezing of gait in parkinson’s disease: a review of two interconnected, episodic phenomena. Movement disorders: official journal of the Movement Disorder Society 19, 871–884 (2004).

[6] Giladi, N. & Nieuwboer, A. Understanding and treating freezing of gait in parkinsonism, proposed working definition, and setting the stage (2008).

[7] Pötter-Nerger, M. & Volkmann, J. Deep brain stimulation for gait and postural symptoms in parkinson’s disease. Movement Disorders 28, 1609–1615 (2013). URL 10.1002/mds.25677.

[8] Castrioto, A. Ten-year outcome of subthalamic stimulation in parkinson disease: A blinded evaluation. Archives of Neurology 68, 1550 (2011). URL 10.1001/archneurol.2011.182.

[9] Oehrn, C. R. et al. Chronic adaptive deep brain stimulation versus conventional stimulation in parkinson’s disease: a blinded randomized feasibility trial. Nature Medicine 30, 3345–3356 (2024). URL 10.1038/s41591-024-03196-z.

[10] Rosin, B. et al. Closed-loop deep brain stimulation is superior in ameliorating parkinsonism. Neuron 72, 370–384 (2011). URL 10.1016/j.neuron.2011.08.023.

[11] Little, S. et al. Adaptive deep brain stimulation in advanced parkinson disease. Annals of Neurology 74, 449–457 (2013). URL 10.1002/ana.23951.

[12] Tinkhauser, G. et al. The modulatory effect of adaptive deep brain stimulation on beta bursts in parkinson’s disease. Brain 140, 1053–1067 (2017). URL 10.1093/brain/awx010.

[13] Feldmann, L. K. et al. Subthalamic beta band suppression reflects effective neuromodulation in chronic recordings. European Journal of Neurology 28, 2372–2377 (2021). URL 10.1111/ene.14801.

[14] Little, S., Pogosyan, A., Kuhn, A. & Brown, P. Beta band stability over time correlates with parkinsonian rigidity and bradykinesia. Experimental Neurology 236, 383–388 (2012). URL 10.1016/j.expneurol.2012.04.024.

[15] Anastasopoulos, D. Tremor in parkinson’s disease may arise from interactions of central rhythms with spinal reflex loop oscillations. Journal of Parkinson’s Disease 10, 383–392 (2020). URL 10.3233/JPD-191715.

[16] McNamara, C. G., Rothwell, M. & Sharott, A. Stable, interactive modulation of neuronal oscillations produced through brain-machine equilibrium. Cell Reports 41, 111616 (2022). URL 10.1016/j.celrep.2022.111616.

[17] Yin, Z. et al. Cortical phase-amplitude coupling is key to the occurrence and treatment of freezing of gait. Brain 145, 2407–2421 (2022). URL 10.1093/brain/awac121.

[18] Toledo, J. B. et al. High beta activity in the subthalamic nucleus and freezing of gait in parkinson’s disease. Neurobiology of Disease 64, 60–65 (2014). URL 10.1016/j.nbd.2013.12.005.

[19] Shine, J. et al. Abnormal patterns of theta frequency oscillations during the temporal evolution of freezing of gait in parkinson’s disease. Clinical Neurophysiology 125, 569–576 (2014). URL 10.1016/j.clinph.2013.09.006.

[20] Neumann, W.-J. et al. Toward electrophysiology-based intelligent adaptive deep brain stimulation for movement disorders. Neurotherapeutics 16, 105–118 (2019). URL 10.1007/s13311-018-00705-0.

[21] Little, S. & Brown, P. What brain signals are suitable for feedback control of deep brain stimulation in parkinson’s disease? Annals of the New York Academy of Sciences 1265, 9–24 (2012). URL 10.1111/j.1749-6632.2012.06650.x.

[22] Wenger, N. et al. Rodent models for gait network disorders in parkinson’s disease – a translational perspective. Experimental Neurology 352, 114011 (2022). URL https://www.sciencedirect.com/science/article/pii/S001448862200036X.

[23] Kiehn, O. Decoding the organization of spinal circuits that control locomotion. Nature Reviews Neuroscience 17, 224–238 (2016). URL 10.1038/nrn.2016.9.

[24] Nishimaru, H., Matsumoto, J., Setogawa, T. & Nishijo, H. Neuronal structures controlling locomotor behavior during active and inactive motor states. Neuroscience Research 189, 83–93 (2023). URL https://www.sciencedirect.com/science/article/pii/S0168010222003017.

[25] Tovote, P. The origins of freezing. Nature Reviews Neuroscience 25, 709–709 (2024). URL 10.1038/s41583-024-00857-3.

[26] Hjorth, B. Eeg analysis based on time domain properties. Electroencephalography and Clinical Neurophysiology 29, 306–310 (1970). URL https://www.sciencedirect.com/science/article/pii/0013469470901434.

[27] Brazhnik, E., Novikov, N., McCoy, A. J., Cruz, A. V. & Walters, J. R. Functional correlates of exaggerated oscillatory activity in basal ganglia output in hemiparkinsonian rats. Experimental Neurology 261, 563–577 (2014). URL https://www.sciencedirect.com/science/article/pii/S0014488614002386.

[28] Xiao, H. et al. Selective cholinergic depletion of pedunculopontine tegmental nucleus aggravates freezing of gait in parkinsonian rats. Neuroscience Letters 659, 92–98 (2017). URL https://www.sciencedirect.com/science/article/pii/S0304394017306638.

[29] Degos, B., Deniau, J.-M., Chavez, M. & Maurice, N. Chronic but not acute dopaminergic transmission interruption promotes a progressive increase in cortical beta frequency synchronization: relationships to vigilance state and akinesia. Cereb. Cortex 19, 1616–1630 (2009).

[30] Merk, T. et al. py neuromodulation: Signal processing and decoding for neural electrophysiological recordings. Journal of Open Source Software 10, 8258 (2025). URL 10.21105/joss.08258.

[31] Urai, A. E., Doiron, B., Leifer, A. M. & Churchland, A. K. Large-scale neural recordings call for new insights to link brain and behavior. Nature Neuroscience 25, 11–19 (2022). URL 10.1038/s41593-021-00980-9.

[32] Schneider, S., Lee, J. H. & Mathis, M. W. Learnable latent embeddings for joint behavioural and neural analysis. Nature 617, 360–368 (2023). URL 10.1038/s41586-023-06031-6.

[33] Barker, M. et al. Introducing the fair principles for research software. Scientific Data 9 (2022). URL 10.1038/s41597-022-01710-x.

[34] Wenger, N. et al. Spatiotemporal neuromodulation therapies engaging muscle synergies improve motor control after spinal cord injury. Nature Medicine 22, 138–145 (2016). URL 10.1038/nm.4025.

[35] Pedregosa, F. et al. Scikit-learn: Machine learning in Python. Journal of Machine Learning Research 12, 2825–2830 (2011).

[36] Li, X., Xing, Y., Martin-Bastida, A., Piccini, P. & Auer, D. P. Patterns of grey matter loss associated with motor subscores in early parkinson’s disease. NeuroImage: Clinical 17, 498–504 (2018). URL 10.1016/j.nicl.2017.11.009.

[37] Salomon, A. et al. A machine learning contest enhances automated freezing of gait detection and reveals time-of-day effects. Nature Communications 15 (2024). URL 10.1038/s41467-024-49027-0.

[38] Dvorani, A. et al. Real-time detection of freezing motions in parkinson’s patients for adaptive gait phase synchronous cueing. Frontiers in Neurology 12 (2021). URL 10.3389/fneur.2021.720516.

[39] Dvorani, A. et al. On-demand gait-synchronous electrical cueing in parkinson’s disease using machine learning and edge computing: A pilot study. IEEE Open Journal of Engineering in Medicine and Biology 5, 306–315 (2024). URL 10.1109/OJEMB.2024.3390562.

[40] Borzì, L., et al. Prediction of freezing of gait in parkinson’s disease using wearables and machine learning. Sensors 21, 614 (2021). URL 10.3390/s21020614.

[41] Koltermann, K. et al. FoG-Finder: Real-time freezing of gait detection and treatment. IEEE Int. Conf. Connect. Health: Appl. Syst. Eng. Technol. 2023, 22–33 (2023).

[42] Pardoel, S. et al. Real-time freezing of gait prediction and detection in parkinson’s disease. Sensors 24, 8211 (2024). URL 10.3390/s24248211.

[43] Wiltschko, A. et al. Mapping sub-second structure in mouse behavior. Neuron 88, 1121–1135 (2015). URL 10.1016/j.neuron.2015.11.031.

[44] Moustafa, A. A. et al. Motor symptoms in parkinson’s disease: A unified framework. Neuroscience & Biobehavioral Reviews 68, 727–740 (2016). URL https://www.sciencedirect.com/science/article/pii/S0149763415300919.

[45] Chaudhuri, K. R., Healy, D. G. & Schapira, A. H. Non-motor symptoms of parkinson’s disease: diagnosis and management. The Lancet Neurology 5, 235–245 (2006). URL 10.1016/S1474-4422(06)70373-8.

[46] Kalia, L. V. & Lang, A. E. Parkinson’s disease. The Lancet 386, 896–912 (2015). URL 10.1016/S0140-6736(14)61393-3.

[47] van Wijk, B. C. M., de Bie, R. M. A. & Beudel, M. A systematic review of local field potential physiomarkers in parkinson’s disease: from clinical correlations to adaptive deep brain stimulation algorithms. Journal of Neurology 270, 1162–1177 (2022). URL 10.1007/s00415-022-11388-1.

[48] Brown, P. Abnormal oscillatory synchronisation in the motor system leads to impaired movement. Current Opinion in Neurobiology 17, 656–664 (2007). URL 10.1016/j.conb.2007.12.001.

[49] Lofredi, R. et al. Beta bursts during continuous movements accompany the velocity decrement in parkinson’s disease patients. Neurobiology of Disease 127, 462–471 (2019). URL 10.1016/j.nbd.2019.03.013.

[50] Litvak, V. et al. Movement-related changes in local and long-range synchronization in parkinson’s disease revealed by simultaneous magnetoencephalography and intracranial recordings. Journal of Neuroscience 32, 10541–10553 (2012). URL 10.1523/JNEUROSCI.0767-12.2012.

[51] Lee, S.-B. et al. Predicting parkinson’s disease using gradient boosting decision tree models with electroencephalography signals. Parkinsonism & Related Disorders 95, 77–85 (2022). URL 10.1016/j.parkreldis.2022.01.011.

[52] Safi, M. S. & Safi, S. M. M. Early detection of alzheimer’s disease from eeg signals using hjorth parameters. Biomedical Signal Processing and Control 65, 102338 (2021). URL 10.1016/j.bspc.2020.102338.

[53] Puri, D. V., Gawande, J. P., Kachare, P. H. & Al-Shourbaji, I. Optimal time-frequency localized wavelet filters for identification of alzheimer’s disease from eeg signals. Cognitive Neurodynamics 19 (2025). URL 10.1007/s11571-024-10198-7.

[54] Coelho, B. F. O. et al. Parkinson’s disease effective biomarkers based on hjorth features improved by machine learning. Expert Systems with Applications 212, 118772 (2023). URL 10.1016/j.eswa.2022.118772.

[55] Yao, L., Brown, P. & Shoaran, M. Improved detection of parkinsonian resting tremor with feature engineering and kalman filtering. Clin. Neurophysiol. 131, 274–284 (2020).

[56] Marquez, J. S. et al. Neural correlates of freezing of gait in parkinson’s disease: an electrophysiology mini-review. Frontiers in neurology 11, 571086 (2020).

[57] Weiss, D. et al. Freezing of gait: understanding the complexity of an enigmatic phenomenon. Brain 143, 14–30 (2020).

[58] Takakusaki, K. Functional neuroanatomy for posture and gait control. Journal of movement disorders 10, 1 (2017).

[59] Pozzi, N. G. et al. Freezing of gait in parkinson’s disease reflects a sudden derangement of locomotor network dynamics. Brain 142, 2037–2050 (2019). URL 10.1093/brain/awz141.

[60] Bharti, K. et al. Neuroimaging advances in parkinson’s disease with freezing of gait: a systematic review. NeuroImage: Clinical 24, 102059 (2019).

[61] Mirelman, A. et al. Gait impairments in parkinson’s disease. The Lancet Neurology 18, 697–708 (2019). URL 10.1016/S1474-4422(19)30044-4.

[62] Revuelta, G. J., Uthayathas, S., Wahlquist, A. E., Factor, S. A. & Papa, S. M. Non-human primate fog develops with advanced parkinsonism induced by mptp treatment. Experimental neurology 237, 464–469 (2012).

[63] Thenaisie, Y. et al. Towards adaptive deep brain stimulation: clinical and technical notes on a novel commercial device for chronic brain sensing. Journal of neural engineering 18, 042002 (2021).

[64] Capogrosso, M. et al. A brain–spine interface alleviating gait deficits after spinal cord injury in primates. Nature 539, 284–288 (2016). URL 10.1038/nature20118.

[65] Lofredi, R. et al. Subthalamic stimulation impairs stopping of ongoing movements. Brain 144, 44–52 (2020). URL 10.1093/brain/awaa341.

