## Supplementary figures for "Deep neurobehavioral phenotyping uncovers neural fingerprints of locomotor deficits in Parkinson’s disease"

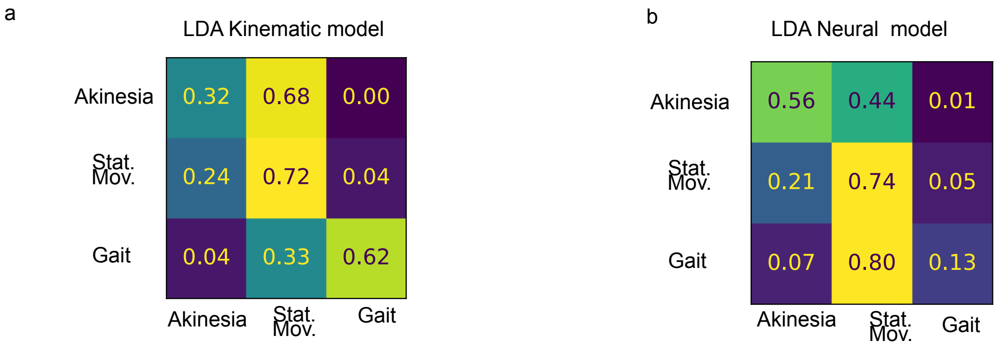


**Fig. 1** **Expanded Confusion Matrixes for LDA and CEBRA models a. LDA kinematic model - Confusion Matrix. b. LDA neural model - Confusion Matrix.**

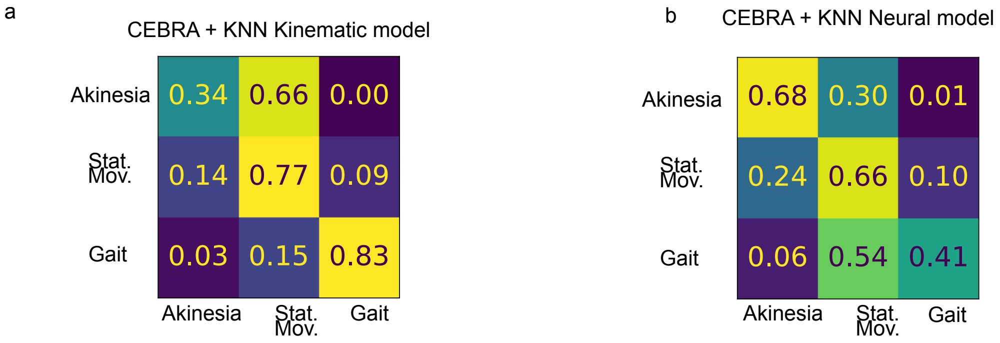


**Fig. 2** **Expanded Confusion Matrixes for CEBRA models + KNN a. CEBRA kinematic model - Confusion Matrix. b. CEBRA neural model - Confusion Matrix.**

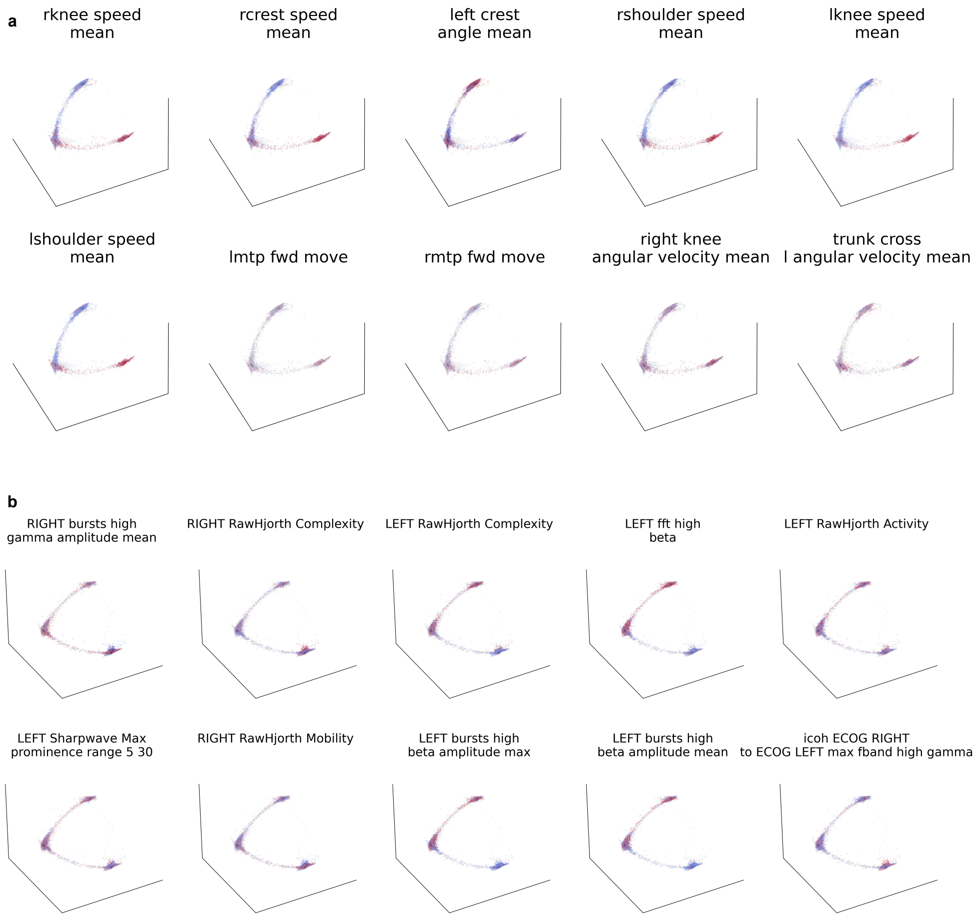


**Fig. 3 Heatmap for top scoring CEBRA features**
**a. Value distribution of top 10 kinematic features in CEBRA kinematic embedding. b. Value distribution of top 10 neural features in CEBRA neural embedding.**

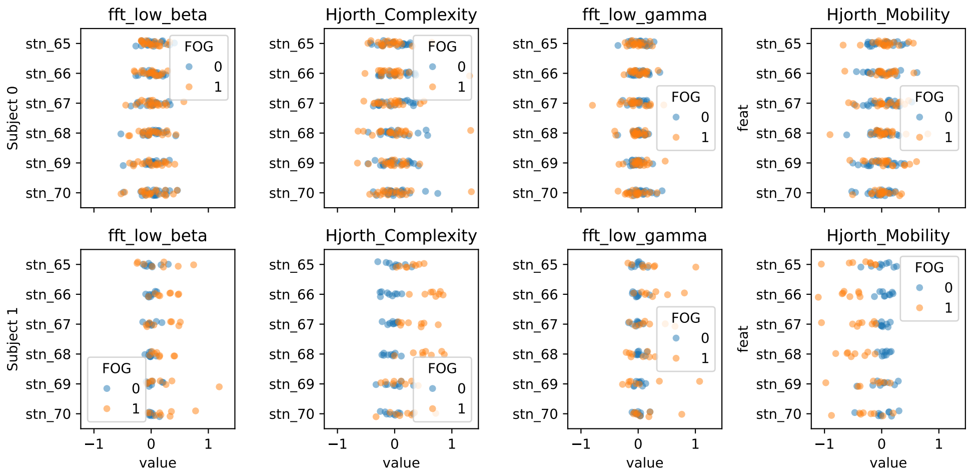


**Fig. 4 Feature distribution in all channels of STN**

| Feature | Description |
|---|---|
| left crest angle mean | Mean angle (shoulder-crest-hip) in window period. Left side. |
| left hip angle mean | Mean angle (crest-hip-knee) in window period. Left side. |
| left knee angle mean | Mean angle (hip-knee-ankle) in window period. Left side. |
| left ankle angle mean | Mean angle (knee-ankle-MTP) in window period. Left side. |
| right crest angle mean | Mean angle (shoulder-crest-hip) in window period. Right side. |
| right hip angle mean | Mean angle (crest-hip-knee) in window period. Right side. |
| right knee angle mean | Mean angle (hip-knee-ankle) in window period. Right side. |
| right ankle angle mean | Mean angle (knee-ankle-MTP) in window period. Right side. |
| trunk cross l angle mean | Mean angle (left shoulder - right crest - left hip) |
| trunk cross r angle mean | Mean angle (right shoulder - left crest - right hip) |
| left crest angular velocity mean | Mean angle velocity (shoulder-crest-hip) in window period. Left side. |
| left hip angular velocity mean | Mean velocity angle (crest-hip-knee) in window period. Left side. |
| left knee angular velocity mean | Mean velocity angle (hip-knee-ankle) in window period. Left side. |
| left ankle angular velocity mean | Mean velocity angle (knee-ankle-MTP) in window period. Left side. |
| right crest angular velocity mean | Mean angle velocity (shoulder-crest-hip) in window period. Right side. |
| right hip angular velocity mean | Mean velocity angle (crest-hip-knee) in window period. Right side. |
| right knee angular velocity mean | Mean velocity angle (hip-knee-ankle) in window period. Right side. |
| right ankle angular velocity mean | Mean velocity angle (knee-ankle-MTP) in window period. Right side. |
| trunk cross l angular velocity mean | Mean angle velocity (left shoulder - right crest - left hip) |
| trunk cross r angular velocity mean | Mean angle velocity (right shoulder - left crest - right hip) |
| lmtp speed mean | Mean speed of MTP. Left side. |
| lankle speed mean | Mean speed of ankle. Left side. |
| lknee speed mean | Mean speed of knee. Left side. |
| lhip speed mean | Mean speed of hip. Left side. |
| lcrest speed mean | Mean speed of crest. Left side. |
| lshoulder speed mean | Mean speed of shoulder. Left side. |

| Feature | Description |
| --- | --- |
| rmtp speed mean | Mean speed of MTP. Right side. |
| rankle speed mean | Mean speed of ankle. Right side. |
| rknee speed mean | Mean speed of knee. Right side. |
| rhip speed mean | Mean speed of hip. Right side. |
| rcrest speed mean | Mean speed of crest. Right side. |
| rshoulder speed mean | Mean speed of shoulder. Right side. |
| lmtp height | Height of MTP. Left side. |
| lmtp fwd move | Relative forward movement of MTP. Left side. |
| rmtp height | Height of MTP. Right side. |
| rmtp fwd move | Relative forward movement of MTP. Right side. |
| ECOG LEFT RawHjorth Activity | Hjorth Activity. Left side. |
| ECOG LEFT RawHjorth Mobility | Hjorth Mobility. Left side. |
| ECOG LEFT RawHjorth Complexity | Hjorth Complexity. Left side. |
| ECOG RIGHT RawHjorth Activity | Hjorth Activity. Right side. |
| ECOG RIGHT RawHjorth Mobility | Hjorth Mobility. Right side. |
| ECOG RIGHT RawHjorth Complexity | Hjorth Complexity. Right side. |
| ECOG LEFT raw | Raw signal. Left side. |
| ECOG RIGHT raw | Raw signal. Right side. |
| ECOG LEFT bandpass activity low beta | Filtered ECoG signal [13, 20] Hz. Left Side. |
| ECOG LEFT bandpass activity high beta | Filtered ECoG signal [20, 35] Hz. Left Side. |
| ECOG LEFT bandpass activity low gamma | Filtered ECoG signal [60, 80] Hz. Left Side. |
| ECOG LEFT bandpass activity high gamma | Filtered ECoG signal [90, 200] Hz. Left Side. |
| ECOG LEFT bandpass activity HFA | Filtered ECoG signal [200, 400] Hz. Left Side. |
| ECOG RIGHT bandpass activity low beta | Filtered ECoG signal [13, 20] Hz. Right Side. |
| ECOG RIGHT bandpass activity high beta | Filtered ECoG signal [20, 35] Hz. Right Side. |
| ECOG RIGHT bandpass activity low gamma | Filtered ECoG signal [60, 80] Hz. Right Side. |
| ECOG RIGHT bandpass activity high gamma | Filtered ECoG signal [90, 200] Hz. Right Side. |
| ECOG RIGHT bandpass activity HFA | Filtered ECoG signal [200, 400] Hz. Right Side. |

| **Feature** | **Description** |
|---|---|
| ECOG LEFT stft low beta | Short-time Fourier transform [13, 20] Hz. Left Side. |
| ECOG LEFT stft high beta | Short-time Fourier transform [20, 35] Hz. Left Side. |
| ECOG LEFT stft low gamma | Short-time Fourier transform [60, 80] Hz. Left Side. |
| ECOG LEFT stft high gamma | Short-time Fourier transform [90, 200] Hz. Left Side. |
| ECOG LEFT stft HFA | Short-time Fourier transform [200, 400] Hz. Left Side. |
| ECOG RIGHT stft low beta | Short-time Fourier transform [13, 20] Hz. Right Side. |
| ECOG RIGHT stft high beta | Short-time Fourier transform [20, 35] Hz. Right Side. |
| ECOG RIGHT stft low gamma | Short-time Fourier transform [60, 80] Hz. Right Side. |
| ECOG RIGHT stft high gamma | Short-time Fourier transform [90, 200] Hz. Right Side. |
| ECOG RIGHT stft HFA | Short-time Fourier transform [200, 400] Hz. Right Side. |
| ECOG LEFT fft low beta | Fast Fourier transform [13, 20] Hz. Left Side. |
| ECOG LEFT fft high beta | Fast Fourier transform [20, 35] Hz. Left Side. |
| ECOG LEFT fft low gamma | Fast Fourier transform [60, 80] Hz. Left Side. |
| ECOG LEFT fft high gamma | Fast Fourier transform [90, 200] Hz. Left Side. |
| ECOG LEFT fft HFA | Fast Fourier transform [200, 400] Hz. Left Side. |
| ECOG RIGHT fft low beta | Fast Fourier transform [13, 20] Hz. Right Side. |
| ECOG RIGHT fft high beta | Fast Fourier transform [20, 35] Hz. Right Side. |
| ECOG RIGHT fft low gamma | Fast Fourier transform [60, 80] Hz. Right Side. |
| ECOG RIGHT fft high gamma | Fast Fourier transform [90, 200] Hz. Right Side. |
| ECOG RIGHT fft HFA | Fast Fourier transform [200, 400] Hz. Right Side. |
| ECOG LEFT Sharpwave Max prominence range 5 80 | Maximum prominence of sharpwave in the 5–80 Hz range. Left Side. |
| ECOG LEFT Sharpwave Mean interval range 5 80 | Mean interval of sharpwave in the 5–80 Hz range. Left Side. |
| ECOG LEFT Sharpwave Max sharpness range 5 80 | Maximum sharpness of sharpwave in the 5–80 Hz range. Left Side. |
| ECOG LEFT Sharpwave Max prominence range 5 30 | Maximum prominence of sharpwave in the 5–30 Hz range. Left Side. |
| ECOG LEFT Sharpwave Mean interval range 5 30 | Mean interval of sharpwave in the 5–30 Hz range. Left Side. |
| ECOG LEFT Sharpwave Max sharpness range 5 30 | Maximum sharpness of sharpwave in the 5–30 Hz range. Left Side. |
| ECOG RIGHT Sharpwave Max prominence range 5 80 | Maximum prominence of sharpwave in the 5–80 Hz range. Right Side. |
| ECOG RIGHT Sharpwave Mean interval range 5 80 | Mean interval of sharpwave in the 5–80 Hz range. Right Side. |

| Feature | Description |
|---|---|
| ECOG RIGHT Sharpwave Max sharpness range 5 80 | Maximum sharpness of sharpwave in the 5–80 Hz range. Right Side. |
| ECOG RIGHT Sharpwave Max prominence range 5 30 | Maximum prominence of sharpwave in the 5–30 Hz range. Right Side. |
| ECOG RIGHT Sharpwave Mean interval range 5 30 | Mean interval of sharpwave in the 5–30 Hz range. Right Side. |
| ECOG RIGHT Sharpwave Max sharpness range 5 30 | Maximum sharpness of sharpwave in the 5–30 Hz range. Right Side. |
| ECOG LEFT fooof a exp | Exponent of the power spectrum model. Left Side. |
| ECOG LEFT fooof a offset | Offset of the power spectrum model. Left Side. |
| ECOG RIGHT fooof a exp | Exponent of the power spectrum model. Right Side. |
| ECOG RIGHT fooof a offset | Offset of the power spectrum model. Right Side. |
| ECOG LEFT bursts low beta duration mean | Mean burst duration in the low beta frequency band (13–20 Hz). Left Side. |
| ECOG LEFT bursts low beta amplitude mean | Mean burst amplitude in the low beta frequency band (13–20 Hz). Left Side. |
| ECOG LEFT bursts low beta duration max | Maximum burst duration in the low beta frequency band (13–20 Hz). Left Side. |
| ECOG LEFT bursts low beta amplitude max | Maximum burst amplitude in the low beta frequency band (13–20 Hz). Left Side. |
| ECOG LEFT bursts low beta burst rate per s | Burst rate per second in the low beta frequency band (13–20 Hz). Left Side. |
| ECOG LEFT bursts high beta duration mean | Mean burst duration in the high beta frequency band (20–35 Hz). Left Side. |
| ECOG LEFT bursts high beta amplitude mean | Mean burst amplitude in the high beta frequency band (20–35 Hz). Left Side. |
| ECOG LEFT bursts high beta duration max | Maximum burst duration in the high beta frequency band (20–35 Hz). Left Side. |
| ECOG LEFT bursts high beta amplitude max | Maximum burst amplitude in the high beta frequency band (20–35 Hz). Left Side. |
| ECOG LEFT bursts high beta burst rate per s | Burst rate per second in the high beta frequency band (20–35 Hz). Left Side. |
| ECOG LEFT bursts low gamma duration mean | Mean burst duration in the low gamma frequency band (60–80 Hz). Left Side. |
| ECOG LEFT bursts low gamma amplitude mean | Mean burst amplitude in the low gamma frequency band (60–80 Hz). Left Side. |
| ECOG LEFT bursts low gamma duration max | Maximum burst duration in the low gamma frequency band (60–80 Hz). Left Side. |
| ECOG LEFT bursts low gamma amplitude max | Maximum burst amplitude in the low gamma frequency band (60–80 Hz). Left Side. |
| ECOG LEFT bursts low gamma burst rate per s | Burst rate per second in the low gamma frequency band (60–80 Hz). Left Side. |
| ECOG LEFT bursts high gamma duration mean | Mean burst duration in the high gamma frequency band (90–200 Hz). Left Side. |

| **Feature** | **Description** |
|---|---|
| ECOG LEFT bursts high gamma amplitude mean | Mean burst amplitude in the high gamma frequency band (90–200 Hz). Left Side. |
| ECOG LEFT bursts high gamma duration max | Maximum burst duration in the high gamma frequency band (90–200 Hz). Left Side. |
| ECOG LEFT bursts high gamma amplitude max | Maximum burst amplitude in the high gamma frequency band (90–200 Hz). Left Side. |
| ECOG LEFT bursts high gamma burst rate per s | Burst rate per second in the high gamma frequency band (90–200 Hz). Left Side. |
| ECOG RIGHT bursts low beta duration mean | Mean burst duration in the low beta frequency band (13–20 Hz). Right Side. |
| ECOG RIGHT bursts low beta amplitude mean | Mean burst amplitude in the low beta frequency band (13–20 Hz). Right Side. |
| ECOG RIGHT bursts low beta duration max | Maximum burst duration in the low beta frequency band (13–20 Hz). Right Side. |
| ECOG RIGHT bursts low beta amplitude max | Maximum burst amplitude in the low beta frequency band (13–20 Hz). Right Side. |
| ECOG RIGHT bursts low beta burst rate per s | Burst rate per second in the low beta frequency band (13–20 Hz). Right Side. |
| ECOG RIGHT bursts high beta duration mean | Mean burst duration in the high beta frequency band (20–35 Hz). Right Side. |
| ECOG RIGHT bursts high beta amplitude mean | Mean burst amplitude in the high beta frequency band (20–35 Hz). Right Side. |
| ECOG RIGHT bursts high beta duration max | Maximum burst duration in the high beta frequency band (20–35 Hz). Right Side. |
| ECOG RIGHT bursts high beta amplitude max | Maximum burst amplitude in the high beta frequency band (20–35 Hz). Right Side. |
| ECOG RIGHT bursts high beta burst rate per s | Burst rate per second in the high beta frequency band (20–35 Hz). Right Side. |
| ECOG RIGHT bursts low gamma duration mean | Mean burst duration in the low gamma frequency band (60–80 Hz). Right Side. |
| ECOG RIGHT bursts low gamma amplitude mean | Mean burst amplitude in the low gamma frequency band (60–80 Hz). Right Side. |
| ECOG RIGHT bursts low gamma duration max | Maximum burst duration in the low gamma frequency band (60–80 Hz). Right Side. |
| ECOG RIGHT bursts low gamma amplitude max | Maximum burst amplitude in the low gamma frequency band (60–80 Hz). Right Side. |
| ECOG RIGHT bursts low gamma burst rate per s | Burst rate per second in the low gamma frequency band (60–80 Hz). Right Side. |
| ECOG RIGHT bursts high gamma duration mean | Mean burst duration in the high gamma frequency band (90–200 Hz). Right Side. |
| ECOG RIGHT bursts high gamma amplitude mean | Mean burst amplitude in the high gamma frequency band (90–200 Hz). Right Side. |
| ECOG RIGHT bursts high gamma duration max | Maximum burst duration in the high gamma frequency band (90–200 Hz). Right Side. |

| **Feature** | **Description** |
|---|---|
| ECOG RIGHT bursts high gamma amplitude max | Maximum burst amplitude in the high gamma frequency band (90–200 Hz). Right Side. |
| ECOG RIGHT bursts high gamma burst rate per s | Burst rate per second in the high gamma frequency band (90–200 Hz). Right Side. |
| coh ECOG RIGHT to ECOG LEFT mean fband low beta | Coherence mean in the low beta frequency band (13–20 Hz) between the right and left sides. |
| coh ECOG RIGHT to ECOG LEFT max fband low beta | Coherence maximum in the low beta frequency band (13–20 Hz) between the right and left sides. |
| coh ECOG RIGHT to ECOG LEFT mean fband high beta | Coherence mean in the high beta frequency band (20–35 Hz) between the right and left sides. |
| coh ECOG RIGHT to ECOG LEFT max fband high beta | Coherence maximum in the high beta frequency band (20–35 Hz) between the right and left sides. |
| coh ECOG RIGHT to ECOG LEFT mean fband high gamma | Coherence maximum in the high gamma frequency band (90–200 Hz) between the right and left sides. |
| coh ECOG RIGHT to ECOG LEFT max fband high gamma | Coherence maximum in the high gamma frequency band (90–200 Hz) between the right and left sides. |
| coh ECOG RIGHT to ECOG LEFT max allfbands high gamma | Coherence maximum across all frequency bands in the high gamma range (90–200 Hz) between the right and left sides. |
| icoh ECOG RIGHT to ECOG LEFT mean fband low beta | Imaginary coherence mean in the low beta frequency band (13–20 Hz) between the right and left sides. |
| icoh ECOG RIGHT to ECOG LEFT max fband low beta | Imaginary coherence maximum in the low beta frequency band (13–20 Hz) between the right and left sides. |
| icoh ECOG RIGHT to ECOG LEFT mean fband high beta | Imaginary coherence mean in the high beta frequency band (20–35 Hz) between the right and left sides. |
| icoh ECOG RIGHT to ECOG LEFT max fband high beta | Imaginary coherence maximum in the high beta frequency band (20–35 Hz) between the right and left sides. |
| icoh ECOG RIGHT to ECOG LEFT mean fband high gamma | Imaginary coherence mean in the high gamma frequency band (90–200 Hz) between the right and left sides. |
| icoh ECOG RIGHT to ECOG LEFT max fband high gamma | Imaginary coherence maximum in the high gamma frequency band (90–200 Hz) between the right and left sides. |
| icoh ECOG RIGHT to ECOG LEFT max allfbands high gamma | Imaginary coherence maximum across all frequency bands in the high gamma range (90–200 Hz) between the right and left sides. |

**Table 1: Full list of features used**
